# Time-dependent chromatin maturation during 3D spheroid culture improves preclinical modeling of non-small cell lung cancer

**DOI:** 10.1101/2025.01.23.634518

**Authors:** Anaïs Darracq, Nicolas Sgarioto, Marielle Huot, Gabrielle McInnes, Antoine Meant, Alexandra Langford, Oscar D Villarreal, François Marois, Maxime Caron, Pascal Saint-Onge, Severine Leclerc, Gaël Cagnone, Gregor Andelfinger, Stéphane Richard, Daniel Sinnett, Serge McGraw, Noël J-M Raynal

## Abstract

Non-small cell lung cancer (NSCLC) is the deadliest cancer worldwide. Therapeutic progress stagnate, highlighting the complexity to replicate NSCLC in preclinical models. Drug discovery studies rely mostly on cancer cells in two-dimension (2D), which poorly predict drug efficacy in patients. There is a growing interest in three-dimensional (3D) preclinical models, such as 3D spheroids, to better model tumor phenotype and improve therapeutic prediction. However, a comprehensive view of 3D culture methods impact on transcriptomes, epigenomes and pharmacological responses and their correlations to NSCLC tumors is still missing. Here, we demonstrate that NSCLC spheroids undergo time-dependent transcriptomic and epigenomic changes, which peak after 3 weeks of culture. While DNA methylome remained stable, chromatin methylation and acetylation marks gained features of advanced NSCLC in a time-dependent manner. Single-cell transcriptomic profiling of spheroids demonstrated that time of 3D culture improved the correlation to NSCLC tumors. Moreover, long-term culture of 3D spheroids increased drug screening predictability, by showing resistance to drugs that failed in NSCLC patients (such as HDAC inhibitors) while demonstrating novel pharmacological vulnerabilities and synergistic interactions (such as combination of PRMT and HDAC inhibitors). Strikingly, reverting 3D spheroids back to 2D culture rapidly reversed transcriptomic, epigenetic and pharmacological signatures acquired after 3 weeks of 3D culture, highlighting the critical impact of cell culture conditions on NSCLC phenotype. Collectively, our findings demonstrate that implementing a time-dependent maturation process into 3D spheroid culture induces chromatin and transcriptomic changes that enhance NSCLC preclinical modeling.

## INTRODUCTION

Non-small cell lung cancer (NSCLC) is the greatest contributor to cancer mortality in the world (∼1.8 million deaths/year).^1^ Although the outcome for early stage NSCLC has improved with the implementation of targeted therapies and immunotherapies over the last decades, the majority of advanced patients develop resistance and ultimately progress.^2^ Thus, advanced NSCLC has still a poor prognosis, with a 5-year survival rate of less than 5% for patients with stage IV disease.^3^ For NSCLC patients lacking targeted therapy option, treatment relies mainly on chemotherapy, which has been the mainstay over the last four decades, with an overall survival of less than two years. The addition of immune checkpoint blockers to platinum-based regimen has improved the overall survival for a sub-group of patients. Despite clinical advances, the median overall survival for metastatic patients still remains low.^4,5^ Thus, there is an urgent medical need for more effective drugs to treat advanced NSCLC.

Frustratingly, only about 5% of new anticancer drugs evaluated in clinical trials obtain regulatory approval, highlighting the limitation of current preclinical models to predict for drug efficacy in patients.^6^ Classical drug discovery pipelines in oncology rely on cancer cell lines, which are typically cultivated in two dimension (2D) plastic dishes.^7,8^ In these artificial conditions, many features of the tumor microenvironment are lacking, including the three dimension (3D) architecture, high cell density, and nutrients and oxygen gradients. In 2D culture, cancer cells exhibit altered transcriptomes, as illustrated by a rapid cell growth.^9,10^ Moreover, 2D culture conditions also impact cancer cell metabolism and consequently cancer epigenome. Indeed, epigenetic marks are directly associated with cell metabolism where S-adenosyl-methionine and acetyl-Coenzyme A are key cofactors derived from metabolism that are used for methylation and acetylation reactions, respectively. Furthermore, high oxygen exposure in 2D culture may also modify cancer cell epigenome because most histone and DNA demethylation reactions are oxygen-dependent.^11^ Thus, 2D culture conditions may alter transcriptome and epigenome of cancer cells, which may create a bias in drug discovery and contribute to the ineffective selection of drug candidates for clinical trials.^7,8,12–14^

Epigenomic marks are dynamically regulated by enzymatic writers that add epigenetic marks onto DNA or histones and enzymatic erasers that remove them. Epigenetic marks regulate chromatin accessibility and thus gene transcription. Open chromatin is associated with gene expression whereas closed chromatin is linked to silencing.^15–17^ The epigenome machinery is a promising target for cancer treatment, as it is rewired dramatically in cancer.^18^ The goal of the epigenetic therapy of cancer aims at inducing tumor suppressor gene reactivation, oncogene silencing, cancer cell differentiation, cell death, and sensitivity to immunotherapy.^3,19,20^ The efficacy of epigenetic drugs has been validated by the approval of DNA methylation, histone acetylation/methylation, and IDH1/2 inhibitors to treat leukemia.^18,21,22^ However, against solid tumors, including NSCLC, all clinical trials to date have failed to corroborate promising preclinical results.^23–25^ Thus, we postulate that preclinical models that better recapitulate NSCLC’s epigenome will help identifying new therapeutic vulnerabilities in NSCLC.

To improve the therapeutic prediction of preclinical models, 3D culture models, such as spheroids, organoids and organ-on-chip have been developed.^10,26–32^ 3D models may help closing the gap between preclinical research and clinical trials, which remains one of the biggest challenges in drug discovery. The rationale is that 3D cell culture may better mimic real tumors, improve the translation of basic and pharmacological studies, and identify more efficient drugs for clinical trials. As compared to organoids and organ-on-chip models, 3D spheroid systems represent accessible and reproducible models, allowing the production of homogeneous spheroids in size and that are scalable for high-throughput drug screening. Generally, 3D spheroids consist of self-assembled spherical aggregates of cancer cells, with high cell density causing nutrient and hypoxia gradients with proliferating cells at the external layer, quiescent cells in the middle and necrotic cells in the inner core.^9^ These features impact drug responses where 3D cancer spheroids show more drug resistance than their 2D cell counterparts.^29,32–38^

However, optimal culture conditions where spheroids most faithfully replicate real tumor transcriptomic, epigenomic and drug sensitivity profiles, remain largely unexplored. We postulated that if cancer cells can retrieve a phenotype that resemble tumors *in situ*, this may occur during a time-dependent process where the 3D microenvironment triggers transcriptomic and epigenomic reprogramming, which can be transmitted throughout several cycle of cell divisions. To define this, we developed a method allowing long-term (>1 month) NSCLC spheroid culture and measured time-dependent transcriptomic and epigenomic changes, specifically on chromatin and at DNA methylation levels. Then, we compared transcriptomic signatures obtained in spheroids to NSCLC patient samples at the single-cell resolution. Using drug screening approaches in monotherapy and combination, we compared the sensitivity of NSCLC 3D spheroids versus cells in 2D. Finally, we studied the plasticity transcriptomic, epigenetic and pharmacological features acquired in 3D when NSCLC are cultivated back into 2D. Altogether, we revealed the occurrence of a time-dependent maturation process associated with epigenetic and transcriptomic reprogramming, which significantly improves the modeling of *in vitro* studies to real NSCLC and reveal novel therapeutic vulnerabilities.

## MATERIALS AND METHODS

### Cell culture in 2D and 3D

The human cell lines A549 (from a white, 58-year-old male with carcinoma; RRID: CVCL_0023) were cultured in F-12K (Kaighn’s Mod.) media (Wisent); Calu-1 (from a 47-year-old, white male patient with epidermoid carcinoma; RRID: CVCL_0608) and Calu-6 (from a 61-year-old, white female patient with anaplastic carcinoma; RRID: CVCL_0236) were cultured in Minimum Essential Medium Eagle’s (EMEM) media (Wisent). Cell culture media were supplemented with 10% Fetal Bovine Serum (FBS, Wisent) and cells were placed in 5% CO_2_ incubator at 37°C. NSCLC cell lines in monolayers (2D) were passaged twice a week to maintain logarithmic phase growth. To form spheroids, 2D cells were trypsinized (Trypsin-EDTA 0.25% with phenol red, Gibco^TM^) and then seeded into Prime Surface 3D Culture Spheroid 96 wells, Ultra-low Attachment (ULA) Plates (SBio®, MS-9096UZ), at a density of 20,000 cells (for A549 and Calu-1 cells) or 10,000 cells (for Calu-6) per well in a volume of 200 µl. ULA plates were centrifuged at 1000 rpm for 5 min to allow the formation of spheroids. After 3 days of 3D culture, spheroids were harvested, pooled together and trypsinized for 5 min at 37°C. Single cells in suspension were seeded back into ULA plates at the initial density and were centrifuged at 1000 rpm for 5 min. This spheroid passage step was then repeated weekly (at day 3, 10, 17, 24… of 3D culture). 50 µL of fresh media were added per well four days after passage. All cell lines were tested regularly for mycoplasma contamination.

### Live-cell imaging

Spheroid compaction was monitored by microscopy using IncuCyte S3 Live-cell imaging system; S3/SX1 G/R optical module (Sartorius). Spheroid images were acquired in phase contrast and bright field with 4X objective. Spheroid areas and diameters were measured using IncuCyte software.

### Viability assay

Cell viability was assessed by flow cytometry. Cells in 3D spheroids and 2D, grown in 96 wells plates were dissociated by trypsinization. Cell culture media was removed prior trypsin addition. Then, trypsinization was stopped by adding the respective media of each well to include floating death cells. Cells were stained with Guava®ViaCount™ reagent (Cytek) according to manufacturer’s guidelines and cell viability was measured on Guava HTS Flow cytometer (Millipore Sigma-Aldrich).

### Cell cycle analyses

Cell suspensions from 2D or 3D cultures were washed with PBS, fixed with 70% cold ethanol and incubated overnight at −20°C. Cells were then washed with PBS and treated with 100 μg/mL RNase A (PureLink RNase A, Invitrogen) for 20 min at 37°C. DNA was stained with 50 μg/mL propidium iodide (Sigma). Cell cycle analysis was performed on Guava HTS Flow cytometer.

### Protein extraction and western blotting

Whole cell extracts were obtained through cell lysis with 25 mM Tris pH 7.3 and 1% sodium dodecyl sulfate. Lysates were then sonicated to release proteins and histones. Total protein concentration was determined using Bradford protein assay. Western blots were performed according to Bio-Rad manufacture’s standard protocol and transfer with polyvinyl difluoride membrane (0.2 µm). The antibodies are listed in Table S1. Images were acquired with Image Quant LAS400 device (GE Healthcare Life Sciences) and bands were quantified with ImageJ software (RRID: SCR_003070).

### RT-qPCR

RNA was extracted with TRIzol (Invitrogen) and treated with DNase I-RNase-free (Thermo Scientific). Reverse transcription reactions were performed using High Capacity cDNA Reverse Transcription Kit (Applied Biosystems). qPCR reactions were done using Taqman Fast qPCR MasterMix (Applied Biosystems) on QuantStudio 7 Flex (Applied Biosystems). qPCR reactions and analysis were carried out by the IRIC Genomic platform, Montreal.

### RNA sequencing and analyses

RNA was extracted from A549 2D and spheroids at different time points using RNeasy Mini kit (QIAGEN) with RNAse treatment according to the manufacturer’s protocol. RNA was then quantified on 2100 Bioanalyzer System (Agilent Technologies). RNA library preparations were carried out on 500 ng of RNA (RIN=10) with TruSeq Stranded Total RNA Sample Preparation Kit (Illumina). The amplified libraries were then purified with beads (AMPure XP, Beckmann Coulter Life sciences) and quantified using the KAPA Universal Library Quantification Kit (KAPA Biosystems - Roche). Sequencing of the libraries was performed on the Illumina NovaSeq 6000 system using 101-bp paired-end sequencing. Paired-end reads were aligned with the Spliced Transcripts Alignment to a Reference (STAR v.2.5.3a) software^39^ on the grch37 (hg19) reference genome.^40^ Gene expression was quantified across all experiments using HTSeq v0.9.0 (RRID: SCR_005514) ^41,42^ with the ENSEMBL gene annotation GRCh37.75. The differential expression relative to the initial time-point was computed using DESeq2.^43^ Significantly upregulated or downregulated genes from day 0 to day 24 were defined according to an absolute log fold change of 1 and a false discovery rate of 0.05. Functional enrichment tests of these genes were performed through the Database for Annotation, Visualization and Integrated Discovery (DAVID) gene ontology enrichment tool (RRID: SCR_001881).^44^

### ChIPmentation sequencing and analyses

Five million A549 cells were cross-linked using 1% formaldehyde. Cells were lysed and the sonication of the nuclei was performed on the BioRuptor UCD- 300 (Diagenode inc.) targeting 150-500 bp size. Immunoprecipitation of the histone marks H3K27ac, H3K4me3, H3K4me1 and H3K36me3 was performed following the Auto-ChIPmentation protocol for Histones (Diagenode inc.) according to the manufacturer’s indications. The libraries were selected by size using Ampure XP Beads (Beckman Coulter) and quantified using the KAPA Library Quantification kit - Universal (KAPA Biosystems). Sequencing of the ChIPmentation libraries was performed on the Illumina NovaSeq 6000 system using 101-bp paired-end sequencing. Paired-end reads were mapped to the hg19 reference genome using the bwa-mem v0.7.12 short read aligner (RRID: SCR_022192).^45–47^ Variable-width peaks were called for broad regions of histone mark enrichment using MACS2 v.2.2.5 and annotated with the Hypergeometric Optimization of Motif Enrichment (HOMER v4.11.1) software (RRID: SCR_010881).^41^ Sets of peaks from the various time-points were merged into a single peak file for each histone mark according to literal overlaps and subsequently ranked by the Spearman correlation between the time-point and the read count of each peak. Heatmaps and average profiles of read coverage across the transcriptional start site (TSS) regions or throughout the gene bodies for each histone mark were plotted with the NGS PLOT software.^48^ Density plots of read coverage across selected genes were made through the Integrative Genomics Viewer (IGV) software.^49^ Selected histone peaks of H3K4me3 and H3K27ac in promoter regions identified transcription factor motifs, which we associated with patient survival. Kaplan-Meier representation were done using cbioportal.org on the Lung Adenocarcinoma dataset.^50^

### Whole genome DNA methylation sequencing and analyses

Genomic DNA extraction was performed using the QIAamp DNA mini Kit (QIAGEN) with RNase treatment according to the manufacturer’s protocol. Genomic DNA was quantified using the Qubit system (ThermoFisher Scientific). Libraries were generated from 1500 ng of genomic DNA spiked with 0.1% (w/w) unmethylated λ DNA (Roche Diagnostics) previously fragmented to 300–400 bp peak sizes using the DMA shearing focused-ultrasonicator E210 (Covaris). Library preparation was done using NxSeq AmpFREE Low DNA Library Kit (Lucigen) according to manufacturer’s instructions, followed by bisulfite conversion with the EZ-DNA Methylation Gold Kit (Zymo Research) according to the manufacturer’s protocol. Libraries were amplified by 6 cycles of PCR using the KAPA Hifi Uracil + DNA polymerase kit (Roche). The amplified libraries were size selected using Ampure XP Beads (Beckman Coulter) and quantified using the KAPA Library Quantification kit (Roche). Sequencing of the WGBS libraries was performed on the Illumina HiSeqX system using 151-bp paired-end sequencing. Analyses were done according to the pipeline established at the McGill Epigenomics Mapping and Data Coordinating Centers.^51,52^ Specific parameters were chosen including 1000 bp step-wise tiling windows, a minimum of 2 CpGs per tile and a minimum of 10× CpG coverage of each tile per sample. The methylation level of a 1000-bp tile was the result of all CpG C/T read counts within the tile. To identify DNA methylation changes, a linear model with the following design: methylation ∼ timepoint was used. Significant DNA methylation changes were designated as tiles with an estimated beta coefficient ± ≥5. HOMERwas used to annotate those sequences.^41^ Gene ontology (GO) terms analyses for RefSeqs associated to promoter-TSS were conducted using cluster Profiler.^53^ To examine the impact of DNA methylation on the regulation of gene expression, Promoter-TSS tiles were mapped to gene expression dataset using UCSC RefGene annotation. Gene Set Enrichment Analysis (GSEA) was conducted on genes with an estimated beta coefficient ± ≥5 for methylation and ± ≥0.5 for expression using Fast Gene Set Enrichment Analysis’s R package.^54^

### Single-cell sequencing analysis

Dissociated single cells A549 were captured in nL droplets using a custom Drop-seq setup according to the standard Drop-seq protocol with minor modifications as described. We used a Medfusion 3500 syringe pumps (Smiths Medical) instead of KD Scientific Legato 100 pumps as well as a different RNase inhibitor (SUPERase In-AM2694; Thermo Fisher Scientific). Single-cell suspensions were run alongside barcoded beads (cell concentration between 120 and 150 cells/µL; bead concentration 140 beads/µL; ChemGenes) to capture single-cell RNA on the barcoded beads. Beads attached to single-cell trancriptomes were subsequently washed, reverse-transcribed then treated with exonuclease.For PCR amplification of single-cell transcriptomes, 4,000 beads (approximately 200 single-cell transcriptomes attached to microparticles) were used as input for each PCR. Individual PCR reactions were subsequently pooled for the desired final number of single-cell transcriptomes attached to microparticles to sequence and cleaned up using AMPure XP (Beckmann Coulter Life Sciences) beads at 0.6 × concentrations twice. Correct size distribution and concentration of complementary DNA was determined using a Bioanalyzer High Sensitivity DNA assay (Agilent Technologies). Library preparation was performed using the Nextera XT DNA Library Preparation Kit (Illumina) with 600 pg input according to the Drop-seq protocol using Illumina i7 primers (N701—N706) together with a custom P5 primer in the library PCR amplification. Libraries were quality-controlled for size and concentration using the Bioanalyzer High Sensitivity DNA assay. Then, libraries were quantified using the Universal Library Quantification Kit (KAPA) before sequencing on a NextSeq 550 system (Illumina) at Génome Québec Integrated Centre for Pediatric Clinical Genomics in Montreal. Sequencing was performed on a NextSeq 550 system with the read settings R1 = 20 bp, R2 = 63 bp and index = 8 bp. Unique molecular identifier (UMI) counts (Drop-seq tool https://github.com/broadinstitute/Drop-seq) for 2D and 3D cultured-cells scRNAseq replicates were merged into one single Digital Gene Expression (DGE) matrix and processed using the Seurat package (Comprehensive Integration of Single-Cell Data) (RRID: SCR_016341).^55^ Cells expressing less than 100 genes and more than 10% of mitochondrial genes were filtered out. Single cell transcriptomes were normalized by dividing by the total number of UMIs per cell, and then multiplied by 10,000. All calculations and data were then performed in log space (i.e. ln(transcripts-per-10,000 +1)). After regressing on the cell cycle, we used the 10 most significant components as input for Uniform Manifold Approximation and Projection (UMAP).

### Comparison of single-cell from A549 cells with single-cell of NSCLC tumours

We selected scRNA-seq publicly available dataset from 18 untreated NSCLC tumors to perform a comparative analysis with A549 cells in 2D and 3D spheroids.^56–60^ Data were visualized and processed using the Seurat package and the R (4.0.0) programming language. Cells expressing less than 200 RNA molecules were filtered out, as well as cells containing more than 10% of mitochondrial RNA. The patient groups were then normalized separately using the SCTransform function in Seurat to homogenise data since different culture and sequencing methods were used for each cohort. UMAPs were generated using the Seurat package (3.1.5) and the R (4.0.0) programming language. The number of principal components (PCs) used to generate the UMAPs was determined graphically with the ElbowPlot function. After calculating the UMAP graphical coordinates with RunUMAP, clusters were determined with the FindNeighbors function, using the most variable genes. Subsequently, the FindClusters function was used with the default settings (resolution = 0.8). In order to establish a parallel between the dataset from patients and A549 cells, we isolated the cancer cells from all the other cell types. For each cohort of patients, Seurat was used to generate lists of the most enriched genes in each cluster with the FindMarkers function using the MAST test. These genes were then compared with a list of markers (Table Suppl. S2) to determine the cell type of each cluster. The non-cancerous cells were then used to perform copy number variability (CNV) analysis and confirm the identity of the cancer cells. The infercnv package (1.4.0) from Bioconductor (RRID: SCR_021140) was used to generate graphs for the visual exclusion of clusters not carrying CNVs. The clusters of cancer cells were isolated in order to perform a projection analysis of them on 2D and 3D culture cells. After isolating the cancer cells for each group, these cells were projected onto the A549 single-cell RNA-seq dataset. UMAP coordinates were calculated for each group and a graphical representation was created by assigning to each cancer cell the most observed culture type (2D, 3D-3d, 3D-10d, and 3D-24d) of neighboring culture cells.

### High-throughput drug screening and drug validation studies

Pharmacological assays were carried out on 2D cells and spheroids (A549 and Calu-6) in 96 well plates. 2D cells were plated 24 hours before the treatments. For drug screening, cells were treated with a library of 181 epigenetic compounds (dissolved in DMSO, L1900 library from Selleckchem) at a dose of 10 µM for 48 hours. Each plate contained internal DMSO control wells. The drug library was also screened in combination with MS023 (Cayman Chemical). In this case, cells were treated simultaneously with 10 µM of the drug library and 100 µM MS023 or DMSO for 48 hours. Cell viability were assessed by flow cytometry using Guava HTS Flow cytometer (Millipore Sigma-Aldrich) using Viacount Staining as previously described.^61^ The percentage of viable cells was calculated and normalized on the internal DMSO controls of each plate. The Synergy Index (SI) was calculated as follows: SI = (Viability in monotherapy × MS023 Viability effect)/Viability in combination.^61^ Independently, the response to MS023 was determined by treating cells with increasing doses of the compound (0.5µM to 150µM) for 48h. GraphPad software (RRID: SCR_002798) was used to generate dose-response curves and calculate IC_50_ values.

## RESULTS

### Long-term 3D spheroid culture protocol for NSCLC

We designed a method allowing long-term 3D spheroid culture of three human *KRAS* driven NSCLC cell lines: A549 (KRAS^G12C^ mutation), Calu-6 (KRAS^Q61K^ mutation) and Calu-1 (KRAS^G12C^ mutation), as *KRAS* mutation is the most common mutation in NSCLC (∼30% of cases).^62^ Cells were expanded in 2D and then trypsinized, before seeding in 96 well ultra-low adhesion plates and assembled by centrifugation to form one single 3D spheroid per well. Optimal seeding concentration per well was determined as the cell density producing the greatest cell viability and growth in 3D; *i.e.* 20,000 cells per spheroid for A549 and Calu-1 cells and 10,000 cells per spheroid for Calu-6 cells. Spheroids were cultivated in the same culture media than their 2D counterparts, without additional growth factors or extracellular matrix. Thus, only one parameter was modified between 2D and 3D cultures (*i.e.* the type of cell culture plates), allowing a direct comparison between conditions. Spheroids were trypsinized and re-seeded at the same density for several weeks to allow time-dependent transcriptomic and epigenetic adaptation to the 3D microenvironment. The dissociation step was done after 3 days of 3D culture and then weekly (on days 10, 17 and 24), which helped maintaining adequate cell viability levels and was compatible for downstream experiments such as transcriptomic, epigenomic and drug screenings (Figure 1a).

**Figure 1.**
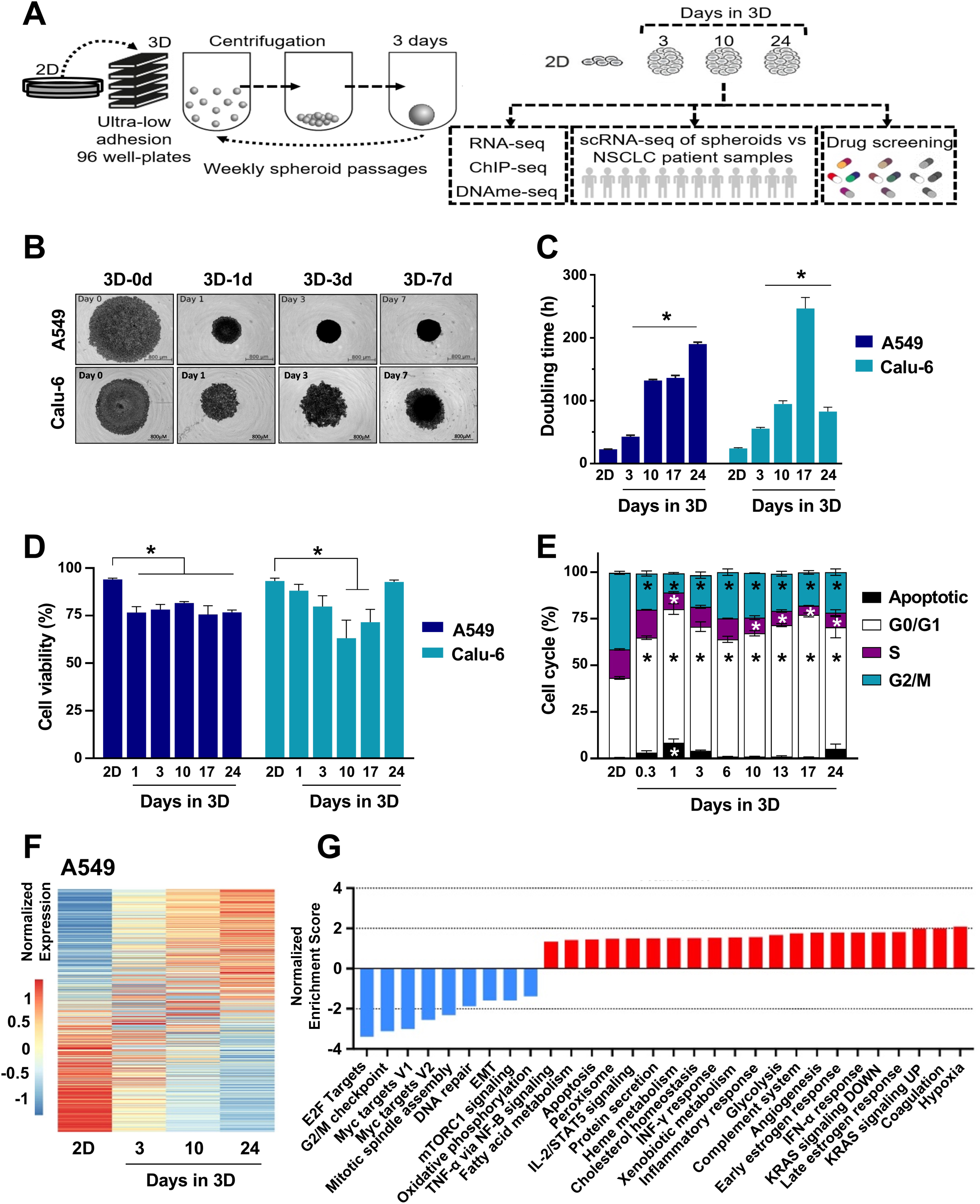
Time-dependent changes in transcriptomic profiles during 3D spheroid culture. **a**) Schematic representation of long-term 3D spheroid formation and experimental design. **b**) Representative bright field microscopy images of A549 and Calu-6 cells in 3D spheroid culture at day 0 (immediately after spheroid assembly), day 1, day 3 and day 7. Pictures were taken with Live Cell Imaging System Incucyte™. Scale bar for all images is 800 µm. **c**) Doubling times (hours) of A549 and Calu-6 cells in 2D and 3D spheroid culture after 3, 10, 17, 24 days. **D**) Cell viability of A549 and Calu-6 cells 3D spheroids after 1, 3, 10, 17, 24 days using cell viability staining (ViaCount™ reagent) relative to cell viability in 2D. **e**) Cell cycle analyses using propidium iodide staining of A549 cells in 2D and 3D culture; days of 3D culture are indicated on the graph. **f**) Heatmap of normalized gene expression levels (25,933 genes) obtained by bulk RNA-sequencing associated in A549 cells in 2D and after 3, 10 and 24 days in 3D spheroids. **g**) Gene set enrichment analysis with normalized enrichment score (NES) for hallmark gene sets associated with the progression from 2D culture method to long-term 3D spheroid culture. Statistically significant changes are indicated by a star (*p* < 0.05 by paired ANOVA).

Within 24 hours after assembly, spheroids of all three cell lines underwent a compaction process in which the diameter of spheroids was reduced by half (less than 1 mm; Figure 1b and Figure S1a-b, Supporting information). Compaction occurred in all spheroids as their diameters were homogeneous within a 96 well-plate (less than 4% variation after 7 days of culture, Figure S1c, Supporting information). Although 3D spheroid diameters were reduced, the number of cells per spheroid increased over time, demonstrating cell growth in 3D (Figure S1d, Supporting information). As expected, cell proliferation rates slowed down significantly during 3D spheroid culture where doubling times of A549 cells reached 132 and 190 hours, after 10 and 24 days of 3D culture, respectively, as compared to 22 hours in 2D. We cultivated spheroids for more than one month, however the doubling times reached 400 hours after 38 days of culture, which made 3D culture less efficient (Figure S1e, Supporting information). Thus, we focused our studies up to the day 24 of 3D culture. Significant increase in doubling times was also observed in Calu-6 spheroids (Figure 1c).

The implementation of dissociation steps was key to obtain long-term 3D culture. The trypsinization at day 3 was instrumental allowing 3D cell growth for more than 10 days but also maintaining healthy 3D culture by removing dead cells accumulated over the first three days of 3D culture. Then, trypsinization was done once a week. Cell viability was measured by flow cytometry. Spheroids had cell viability levels ∼ 75% during 3D culture with A549 cells, which was significantly less than 2D conditions (∼ 90%) but still at levels compatible with drug screenings and other downstream applications (Figure 1d). For Calu-6 spheroids, cell viability decreased significantly on days 10 and 17 (to ∼ 60%) and returned to 93% at day 24, as in 2D condition, which correlated with the doubling time rates (Figure 1c-d). Additionally, Calu-1 spheroid viability decreased over the time and stabilized at ∼ 50%, on days 17 and 24 (Figure S1f, Supporting information). Cell cycle analysis showed that A549 spheroids were characterized by a greater proportion of cells in G0/G1 phase and a reduction in S and G2/M phases, supporting the decrease in proliferation rate observed during 3D culture. In addition, spheroids showed an increased apoptosis during the first days of culture, supporting the dissociation step implemented to the protocol on day 3 (Figure 1e).

To define whether 3D culture had an immediate, a delayed or time-dependent effect on gene expression, we used bulk RNA-Seq on A549 cells in 2D and 3D spheroid culture after 3, 10 and 24 days. Transcriptomic profiles showed a transcriptomic shift from 2D condition throughout the 3D spheroid culture, demonstrating a time-dependent transcriptomic change (Figure 1f and Figure S2a, Supporting information). Transcriptomic profiles showed a clear separation between cell culture conditions on heat map and principal component analyses (Figure S2b-d, Supporting information). Genes that gained the greatest expression in 3D spheroids (at day 24) are markers of lung cancer tissues such as *CST1*, *CEACAM5*, and *CEACAM6* (Figure S2c, Supporting information). For instance, high expression of CEACAM5 is reported in 20% of NSCLC clinical samples, and CST1 is considered as diagnostic marker for NSCLC.^63,64^ Moreover, we found *LEF1*, *ERBB2* and *IL2* as the most oncogenic gene set enriched in 3D and *CSR*, *RB* and *RPS14* as the most downregulated gene sets. 3D culture was also associated with an enrichment of carbohydrate transmembrane transporter and hydrolase activity, suggesting significant metabolic changes (Figure S2d-e, Supporting information). Since transcriptomic pathways were differentially modified in a time-dependent manner during 3D culture, we expressed, by linear regression, a normalized enrichment score for pathway analysis from 2D conditions to different times of 3D spheroid culture. Notably, transcriptomic pathways were enriched in hypoxia, KRAS signaling, glycolysis, inflammatory response, during 3D culture, while a reduction in G2/M checkpoint, Myc targets, DNA repair and epithelial and mesenchymal transition (EMT) pathways was observed (Figure 1g and Figure S3a-d, Supporting information). Reduction of EMT phenotype was validated by RT-qPCR with *E-Cadherin*, *Slug* and *Snail* downregulation, while *N-Cadherin* was upregulated (Figure S3e-f, Supporting information). In agreement with these results, A549 cells from 3D spheroid cultures after 24 days showed a significant reduction in invasion by 75% as compared to A549 cells in 2D cultures (Figure S3g, Supporting information). Altogether, we showed that long-term 3D culturing of NSCLC cells triggers time-dependent transcriptomic changes.

### Time-dependent changes in epigenetic regulator levels during 3D spheroid culture

To investigate whether the epigenome could be altered over the course of 3D culture, we first investigated whether the expression of epigenetic related genes was impacted by 3D culture conditions over time. Using our bulk RNA-seq dataset, we identified 75 epigenetic genes (involving epigenetic writers and chromatin remodelers) showing significant differential mRNA expression between 2D and 3D cultures in A549 cells (Figure 2a). To further assess the expression changes of epigenetic regulators in A549 3D spheroids, we looked at protein expression of a series of epigenetic writers, erasers, chromatin remodelers and histone modifications in various spheroids lines (from A549, Calu-6 and Calu-1 cells). In A549 spheroids, we measured a significant downregulation of acetyltransferases (KAT2A, KAT3A/B, KAT7) and an upregulation of KAT6A and HDAC6 protein levels during 3D culture as compared to the parental cells in 2D (Figure 2b). Interestingly, HDAC6 overexpression is correlated with tumorigenesis.^65^ No significant difference was observed for other histone deacetylase (HDAC) family members (Figure S4a, Supporting information). The remodeler BRG1 (also known as SMARCA4), the catalytic subunit of the SWI/SNF complex and the demethylase LSD1 (Lysine-specific histone demethylase 1A) were both significantly downregulated in A549 spheroids in a time-dependent manner. Interestingly, BRG1 plays a critical role in NSCLC, since its inactivation promotes cancer aggressitivity through chromatin remodeling.^66,67^ Several histone modifications were modified, including a significant increase of the activating marks H3K9ac, H4K16ac, H4R3me2a (histone 4 arginine 3 asymmetric dimethylation) and the repressive marks H3K9me (mono-, di- and tri- methylation), H3K27me3 during the 3D culture, as compared to the 2D condition (Figure 2d and Figure S4b-d, Supporting information). Protein expression of epigenetic regulators was also modified in Calu-6 and Calu-1 spheroids. KAT6A, the methyltransferase G9a and the demethylase LSD1 were upregulated in Calu-6 spheroids. Interestingly, G9a expression was reported to promote NSCLC growth.^68^ In addition, the marks H3K9me1, H3K27me3 and H4R3me2a were significantly increased (Figure 2e). Whilst Calu-1 spheroids showed KAT6A, HDAC6 and EZH2 downregulation, the H4R3me2a mark showed a significant time-dependent increase as in the other spheroid models (Figure 2f-g). Interestingly, type I protein arginine methytransferase 1 (PRMT1), which catalyzes H4R3me2a mark, is key therapeutic target in NSCLC.^69–71^ Overall, long-term 3D culture is characterized by time-dependent changes in the expression of epigenetic proteins resulting in global modifications of specific post-translational histone modifications, such as H4R3me2a.

**Figure 2.**
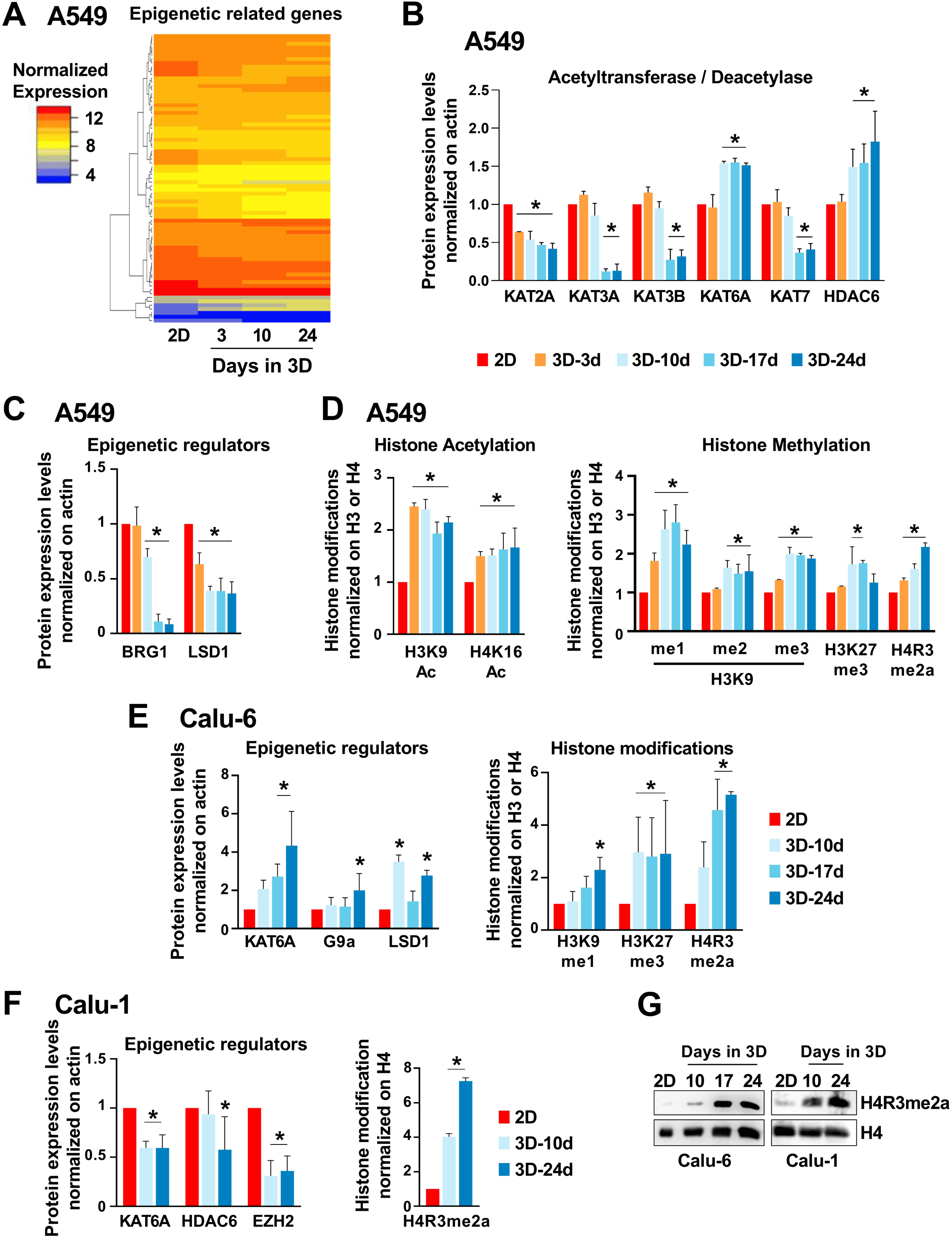
Time-dependent changes of epigenetic regulator expression levels during 3D spheroid culture. **A**) Heatmap clustering of RNA-seq values in 75 curated genes (with significant *p* values) involved in epigenetic regulation in A549 cells in 2D and 3D spheroids. Clustering was performed using Heatmapper.ca of normalized gene expression levels (average linkage clustering method with Euclidian with distance measurement method). **b-c-e-f)** Quantification of protein expression levels in 3D spheroids (3, 10, 17, 24 days) relative to 2D condition in A549 (**b-c**); Calu-6 (**e**, left panel) and Calu-1 (**f**, left panel) cell lines (mean ± SEM, n = 3; a star indicates *p* < 0.05 by ANOVA). ACTIN was used as loading control. **d-e-f**) Quantification of post-translational modifications on histone H3 and H4 proteins of A549 (**d**), Calu-6 (**e**, right panel), Calu-1 (**f**, right panel) in 3D relative to 2D (mean ± SEM, n = 3; a star indicates *p* < 0.05 by ANOVA). Total H3 and H4 were used as loading controls, for H3 and H4 modifications, respectively. **g**) Representative western blotting images of H4R3me2a and H4 proteins in Calu-6, Calu-1, in 2D cultures and 3D spheroids.

### 3D culture triggered time-dependent chromatin changes whereas DNA methylation was not impacted

To provide a comprehensive view of the epigenetic changes at the genomic level, we investigated chromatin post-translational modifications by ChIPmentation and DNA methylation by whole genome bisulfite sequencing in A549 cells in 2D and after 3, 10 and 24 days of spheroid culture. First, we determined the enrichment of active histone marks such as H3K4me1, H3K4me3, H3K27ac and H3K36me3. For H3K4me3 and H3K27ac marks, most changes occurred at the transcriptional start site (TSS), where we observed a reduction in H3K4me3, H3K4me1 and H3K27ac peaks starting rapidly from day 3 of 3D culture and increasing overtime (Figure 3a-b and Figure S5a, Supporting information). H3K27ac decrease occurred more rapidly than H3K4me3. Similarly, gene body regions were marked by a global decrease in H3K4me1 and H3K36me3 (Figure S5a-b, Supporting information). Notably, no significant variation in the total quantity of these histone marks was detected by Western blotting (Figure S4b, Supporting information and data not shown), suggesting the redistribution of theses marks throughout the genome during 3D culture. A reduction in the activating marks, H3K4me3 and H3K27ac, in TSS regions resulted in loss of gene expression, as exemplified by the silencing of *CDH4* gene (Figure 3c).

**Figure 3.**
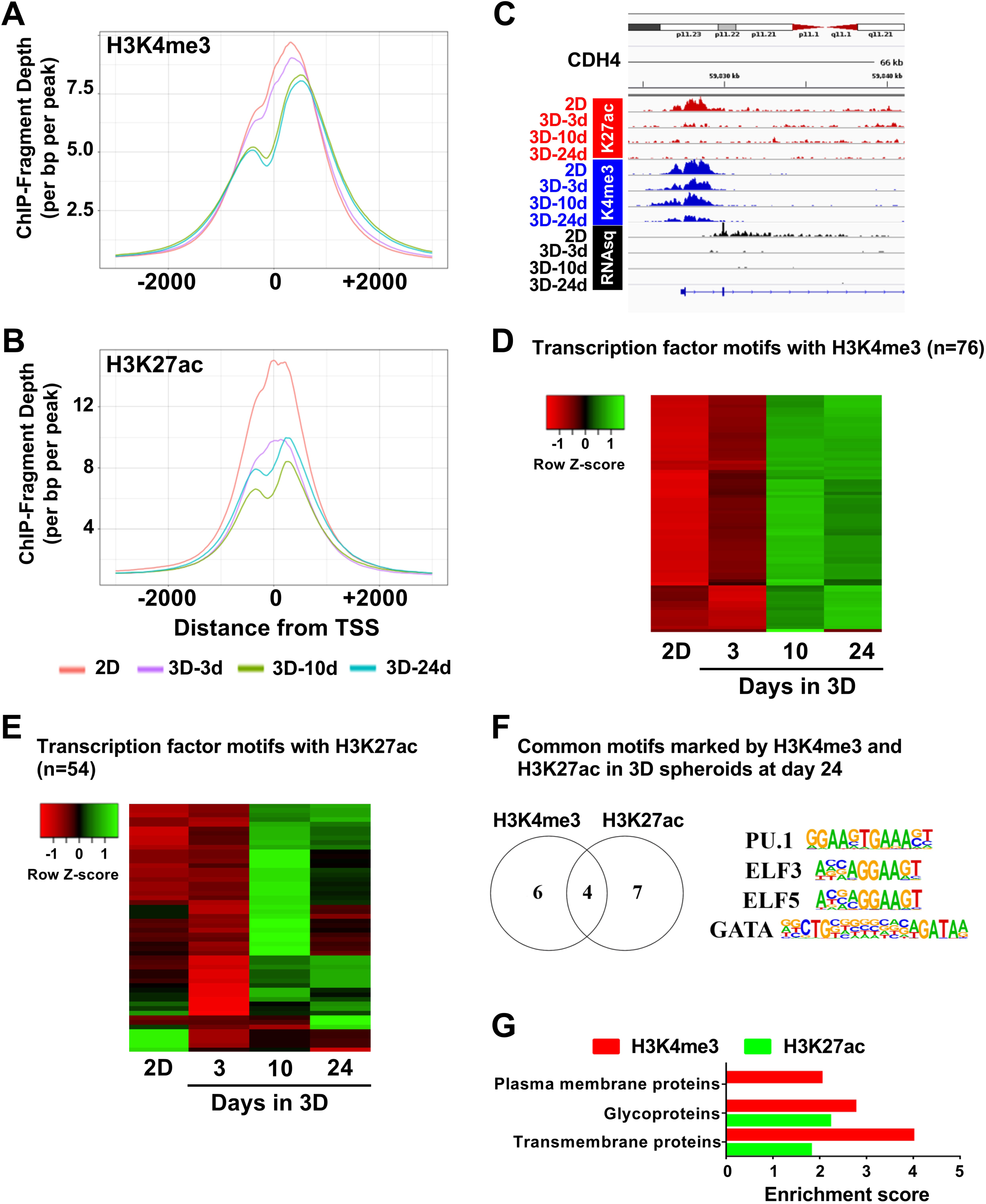
Chromatin changes during 3D culture over time. **a-b**) Sequencing chromatin modifications for H3K4me3 (**a**) and H3K27Ac (**b**) marks centered on transcription start sites (TSS) in A549 cells in 2D and in 3D spheroids after 3, 10 and 24 days. **c**) Visualization of H3K27ac, H3K4me3 and RNA-sequencing enrichments of the CDH4 gene by Integrative Genomics Viewer in A549 cells in 2D and spheroids after 3, 10 and 24 days. **d-e**) Heatmap shows gene expression changes of transcription factor motifs marked by H3K4me3 (n = 76) (**d**) and H3K27ac (n = 54) (**e**) peak enrichment (identified by Hypergeometric Optimization of Motif EnRichment-HOMER) on A549 cell culture in 2D and in spheroids after 3, 10 and 24 days. **f**) Venn diagram shows common motifs marked by H3K4me3 and H3K27ac in 3D spheroids of A549 cells at day 24. **g**) Enrichment scores in pathways of gene promoters marked by H3K4me3 and H3K27ac.

Using HOMER software, we asked whether specific transcription factor binding motifs are associated with gain or loss of H3K4me3 and H3K27ac peaks, in promoter regions, during 3D culture. We identified a series of 76 and 54 transcription factor motifs that mostly gained H3K4me3 and H3K27ac peaks during 3D culture of A549 spheroids (Figure 3d-e, and Table S3, Supporting information). We then focused on the common motifs marked by H3K4me3 and H3K27ac in promoter regions after 24 days of 3D culture. We identified 4 binding motifs of *PU.1*, *ELF3*, *ELF5* and *GATA* transcription factors (Figure 3f). Interestingly, high expression of *ELF3*, and *ELF5* are associated with a significant lower survival rate in NCSLC patients, demonstrating that time-dependent changes induced by 3D culture produce chromatin and transcriptional states associated with worse survival (Figure S5, Supporting information).^72,73^ Lastly, we asked what gene ontology pathways are associated with gains in H3K4me3 and H3K27ac marks. These activating marks in promoter regions are associated with an enrichment several pathways involving glycoproteins, transmembrane proteins, plasma membrane proteins, and KRAS signaling (Figure 3g and Figure S6a-b, Supporting information). To assess whether DNA methylation levels were impacted during 3D culture of A549 spheroids, we performed whole genome bisulfite sequencing and correlated DNA methylation with bulk transcriptomic data. The analyses showed no significant correlation between DNA methylation variations with gene expression at promoter and intragenic regions (Figure S7a-c, Supporting information). Altogether, the results support that 3D culture produces time-dependent chromatin changes at the genomic scale, which peaked after 24 days of spheroid culture.

### Single-cell RNA (scRNA)-sequencing reveals a marked transcriptomic evolution from 2D culture to long-term 3D culture

We next performed scRNA-seq to follow the evolution of the transcriptomic signature and to evaluate its heterogeneity over the time throughout 3D culture in A549 cells. To obtain a global representation of the dataset, we applied the non-linear dimensionality reduction visualization algorithm, Uniform Manifold Approximation and Projection (UMAP; Figure 4a, Supporting information). Single-cell transcriptomic analysis demonstrated a clear separation between 2D cells and 3D spheroids at different time points (days 3, 10 and 24). Transcriptomes after 3 days of 3D culture were clearly separated from those of 2D cells, highlighting the rapid transcriptional reprogramming. Similarly, single-cell transcriptomes of 3D spheroids clustered separately for all 3 time points, supporting both bulk RNA-seq (Figure 1f) and epigenetic changes (Figure 2b-g). Using a clustering analysis, we identified 15 clusters among all cell populations (from 2D to 3D spheroids) where clusters 1 to 4 corresponded to 2D cells, clusters 5 to 8 corresponded to spheroids at day 3, clusters 9 to 11 corresponded to spheroids at day 10, and clusters 12 to 15 corresponded to spheroids at day 24 (Figure 4b). To represent the clustering analysis, we showed the proportion of A549 cells from the different culture condition within each individual cluster (Figure 4c). The data reveal a clear separation of each cell culture condition among the 15 clusters. Then, we performed single-cell gene ontology analysis among the 15 clusters. Among the most differentially expressed gene set, MYC targets (V1 and V2) and oxidative phosphorylation pathways were expressed in 2D and downregulated in spheroids from 10 days of 3D culture (Figure 4d). Conversely, hypoxia, inflammatory pathways and KRAS signaling pathways became expressed in spheroids after 10 and 24 days of 3D culture, further supporting a time-dependent impact during 3D culture where a delayed activation is observed for some key gene pathways (Figure S8a, Supporting information). Then, we plotted the most differentially expressed genes among the 15 different clusters. *CST4*, *CST1* and *CEACAM6* were highly expressed in cluster 15 (belonging to spheroids at 24 days) and lacking in 2D, while caveolin 1 (CAV1) was only found expressed in 2D cells (Figure 4e). The genes were also identified by bulk RNA-seq (Figure S2c, Supporting information). Interestingly, *CST1* and *CST4* are members of type II cystatins protein family, involved in tumor progression and metastasis in different types of cancers including lung cancer. CST1 expression in NSCLC samples is associated with a poor prognosis.^64^ CEACAM6 is overexpressed in most lung adenocarcinomas and is associated with poor overall survival in patients.^74^ We confirmed at the protein level, that CST1, CST4 and CEACAM6 were significantly expressed in a time-dependent manner in spheroids whereas CAV1 was silenced (Figure 4f). Overall, scRNA-seq data refined our bulk RNA-seq results, highlighting a significant transcriptional shift, with clear separation between cell populations produced by long-term 3D culture. The presence of differentially expressed markers show that 3D spheroids more closely resemble NSCLC than cells cultured in 2D.

**Figure 4.**
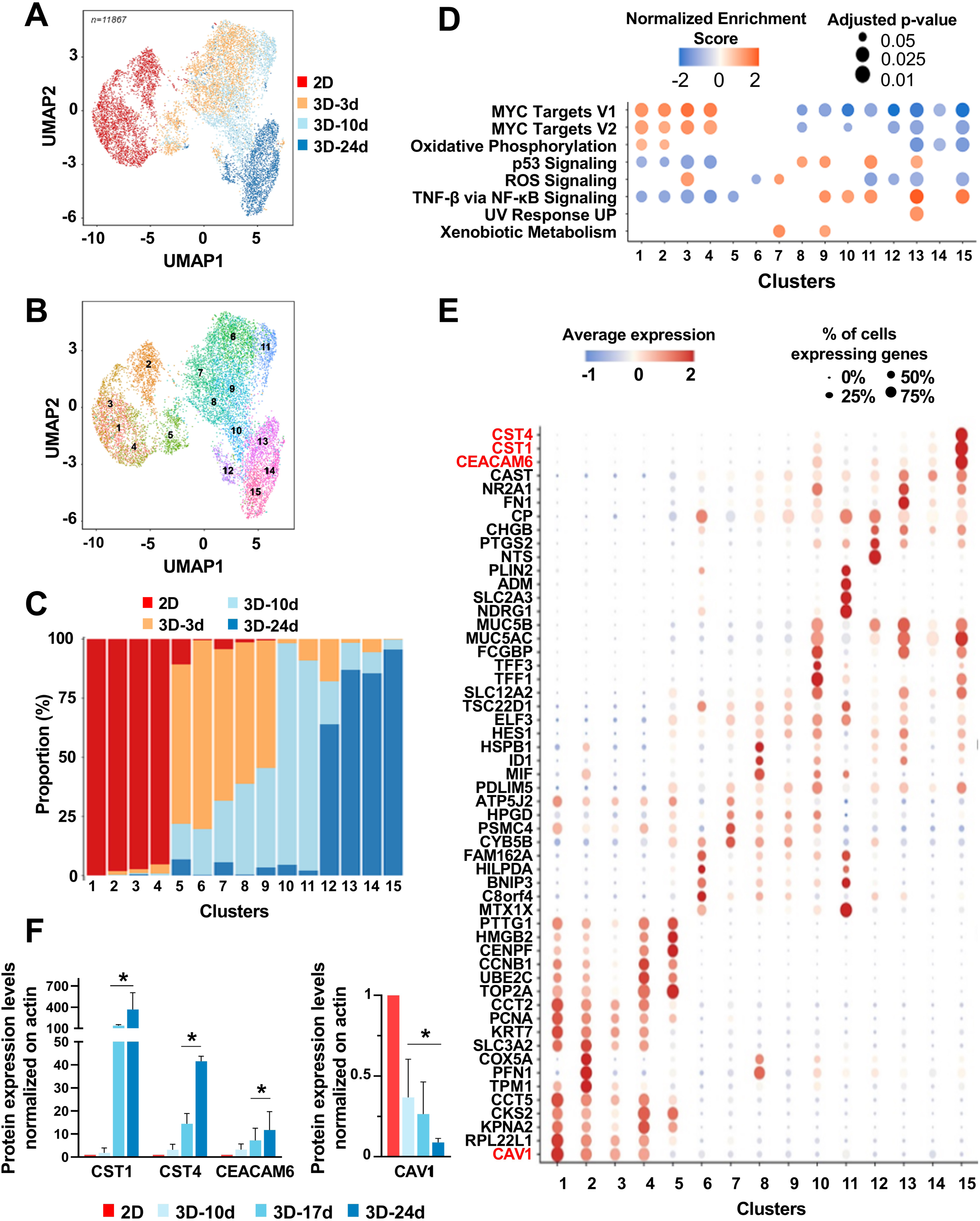
Single-cell RNA sequencing evolution from 2D culture to long-term 3D culture. **a**) Uniform Manifold Approximation and Projection (UMAP) of single-cell RNA sequencing (scRNA-seq) of A549 cells (n = 11,867) in 2D and in 3D after 3, 10 and 24 days. **b**) Seurat clustering analyses of scRNA-sequencing. **c**) Relative proportions of 2D cells and 3D spheroids at different time points in the 15 clusters, identified in **b**. **d**) Single-sample gene set enrichment analysis between the 15 different clusters. Additional pathway enrichment is shown on Supplementary figure S9. Adjusted *p*-value and normalized enrichment score scales are shown in the graph. **e**) Most differentially expressed genes among the 15 clusters identified by Seurat. Average gene expression is shown in blue-grey colors (log2 fold-change). Circle size shows the percent of cells expressing the gene within the cluster. **f**) Quantification of protein expression levels in A549 spheroids (10, 17, 24 days) relative to 2D condition (mean ± SEM, n = 3; *p* < 0.05 by ANOVA). Actin was used as loading control.

### Transcriptomes of spheroids after 24 days of 3D culture better resemble those of NSCLC tumors

To develop preclinical models with enhanced predictive values for drug screening and drug efficacy evaluation, cancer cells in 3D spheroids should better mimic transcriptomic profiles of NSCLC tumors, as compared to classical models. To address this important question, we compared the scRNA-Seq from A549 cells in 2D and in 3D spheroids at the 3 time points (day 3, 10, and 24) to publicly available scRNA-Seq dataset from 18 untreated NSCLC primary tumor samples (to avoid the impact of therapy to gene expression profiles, Figure 5a).^56–60^ We showed a UMAP representation of the 18 tumor patient samples (Figure 5b), which were composed of 8 different cell types (natural killer cells, B cells, T cells, myeloid cells, mastocytes, endothelial cells, fibroblasts and malignant cells) that we annotated based on the expression of specific cell population markers (Figure 5c). The cellular heterogeneity of NSCLC tumors was consistent with the NSCLC inter-patient variability observed in clinic (Figure S8b, Supporting information). To validate our cell-annotation, we compared the single-cell transcriptomic profiles of the 8 different cell types to the A549 in the different cell culture conditions. A Spearman correlation showed that A549 cells when cultured in spheroids after 10 and 24 days showed a better correlation to the malignant cells of the 18 different patients, supporting our cell-annotation (Figure S8c, Supporting information). Then, the transcriptomes of isolated malignant cells from the 18 NSCLC patients (6,856 cells) were compared to A549 cells in 2D and in spheroids (at day 3, 10 and 24; a total of 11,867 cells). We used a projection experiment to visualize the correlation between patient samples and A549 cells in the different cell culture conditions. First, we generated a UMAP representation of A549 cells with the 2D and 3D conditions (Figure 5d, left panel) and a UMAP representation where the A549 scRNA-seq are colored in grey and where tumor cells from patients were projected on the UMAP (Figure 5d, right panel). Interestingly, tumor transcriptomic signatures from the 18 patients matched 99% with A549 3D spheroids, whereas minimal transcriptomic match was observed with A549 cells in 2D. These findings suggest that 2D cultures do not mimic the transcriptome of NSCLC patients. Few patient cells clustered with A549 spheroids after 3 and 10 days of 3D culture. However, the largest transcriptomic correlation was observed between the patient samples and the A549 after 24 days of 3D culture. Strikingly, when we calculated the relative proportion of A549 cells in 2D that mimicked the transcription of the 18 patient samples, it was negligible, supporting the lack of tumor representation of 2D culture (Figure 5f). Importantly, all patient samples, except for patient 1, showed an increased transcriptomic correlation with 3D spheroids at days 3 and 10 but with the greatest proportion of 50% and more at day 24, supporting the time-dependent maturation during 3D spheroid culture. Lastly, Spearman correlation confirmed the improved transcriptomic mimicry of A549 cells in spheroids to the patient samples (Figure 5f). Altogether, these results demonstrate that spheroids undergo a time-dependent maturation process, which better mimic NSCLC patient samples, suggesting that long-term spheroids may represent better preclinical models for drug efficacy studies.

**Figure 5.**
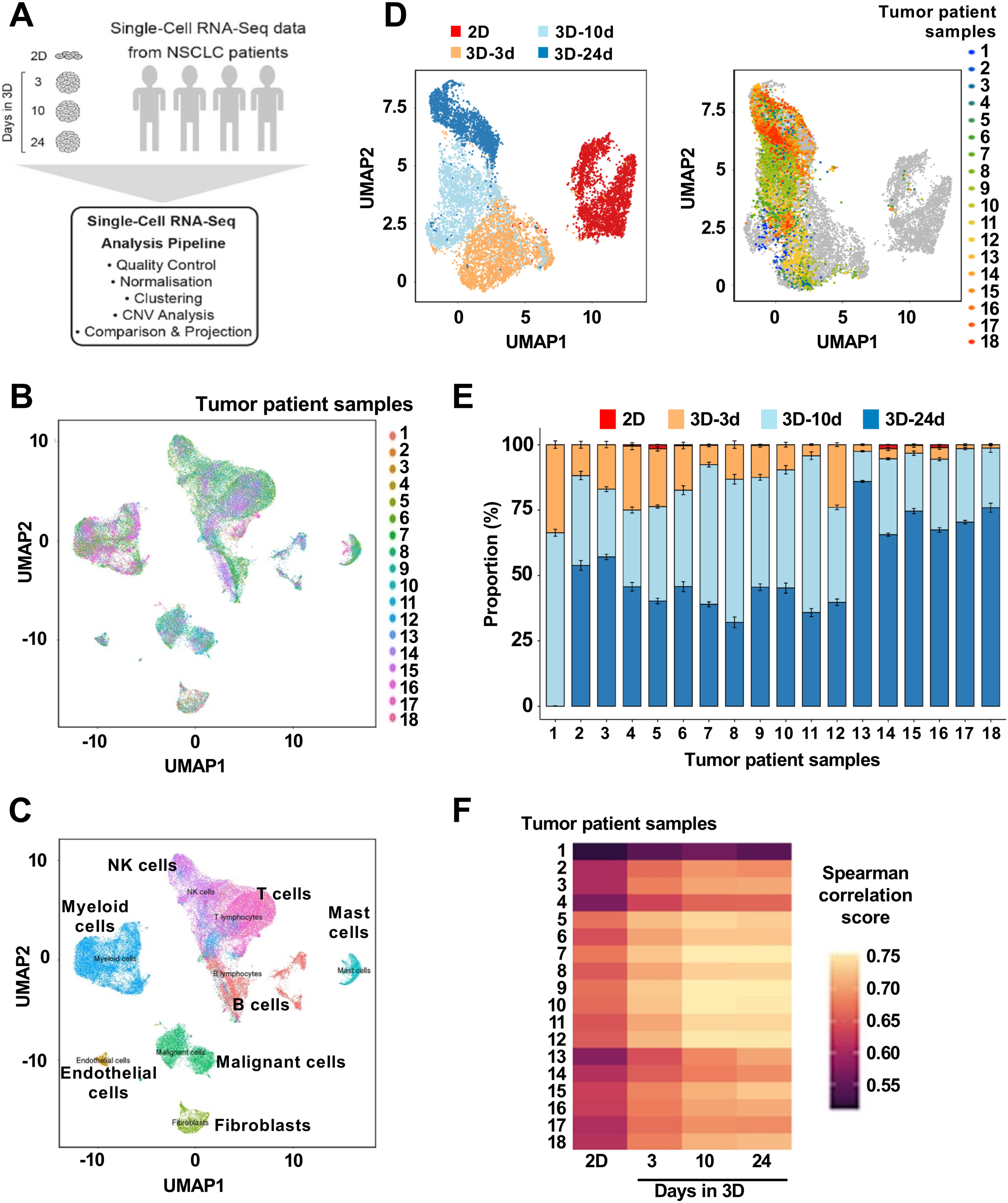
Transcriptomes of 3D spheroids closely resemble those of NSCLC tumors. **a**) Schematic of scRNA-seq pipeline comparing transcriptomic profiles of A549 cells and 18 untreated NSCLC patient samples. **b**) UMAP of scRNA-seq of the 18 NSCLC patients. Each patient’s cell is color coded. **c**) UMAP of scRNA-seq of different cell populations identified by cell markers among the 18 untreated NSCLC patients. Malignant cells are shown in green (n = 6,856). **d**) Left panel, UMAP of A549 cells scRNA-seq data in 2D and 3D after 3, 10 and 24 days of culture (11,867 cells). Each dot represents a single cell from A549 cell population. Right panel, scRNA-seq projection of 18 NSCLC samples (6,856 cells) on A549 cells in 2D and 3D spheroids at different time in culture. Each color dot represents a single tumor cell from a given patient (n = 18). Each grey dot represents cells from A549 cells. **e**) Proportion of single cells within each tumor sample (n = 18) whose transcriptome match those of A549 cells in different conditions (2D, 3D for 3, 10 and 24 days). Means and standard deviations were calculated from 10 projection runs. **f**) Spearman correlation scores between A549 cells (in 2D and in spheroids at 3, 10, and 24 days) versus tumor cells isolated in the 18 NSCLC tumor samples.

### Drug screening with long-term 3D spheroid culture reveals new therapeutic vulnerability

We tested our long-term spheroid preclinical models in drug screening. A549 and Calu-6 cells in 2D and 3D spheroids (after 24 days) were treated with an epigenetic drug library of 181 compounds for 48 hours, at 10 µM and we measured cell viability by flow cytometry (Figure 6a). As expected, 3D spheroids were generally more resistant than their 2D counterparts (Figure 6b and Figure S9a-b, Supporting information). Drug diffusion within spheroids was validated in independent experiments, which pointed towards intrinsic drug resistance rather than drug diffusion issues (data not shown). The drug library is composed by 15 drug target classes, which allowed to highlight class effects. For instance, A549 cells in 2D showed therapeutic vulnerabilities to FLT3 and HDAC inhibitors. In contrast, A549 spheroids became generally more resistant to all drugs including FLT3 and HDAC inhibitors. (Figure S9c, Supporting information). Similar results were observed with Calu-6 cells, although few HDAC inhibitors remained active (Figure S9c, Supporting information). It is noteworthy that HDAC inhibitors showed promising anticancer effects against many lung cancer preclinical studies but failed to demonstrate similar efficacy in patients. Thus, we focused on this drug class in more details, as a typical example of compounds showing efficacy in the laboratory but lacking efficacy in patients. The epigenetic library contains 32 HDAC inhibitors: 18 of those induced at least 50% cell death in 2D and spheroids gained resistance to those HDAC inhibitors (Figure 6c, Supporting information). Thirteen of them showed no efficacy in 2D, which was similar in 3D. Only MC1568 maintained its efficacy in both 2D and 3D models. Similarly, Calu-6 cells gained resistance to some HDAC inhibitors in spheroids compared to 2D. However, 5 HDAC inhibitors were more potent by inducing more than 10% cell death in spheroids, as compared to 2D (Figure 6c, Supporting information). Altogether, the results show that long-term spheroids are more stringent *in vitro* models than classical 2D cells that can better predict some drug failure in clinical trials.

**Figure 6.**
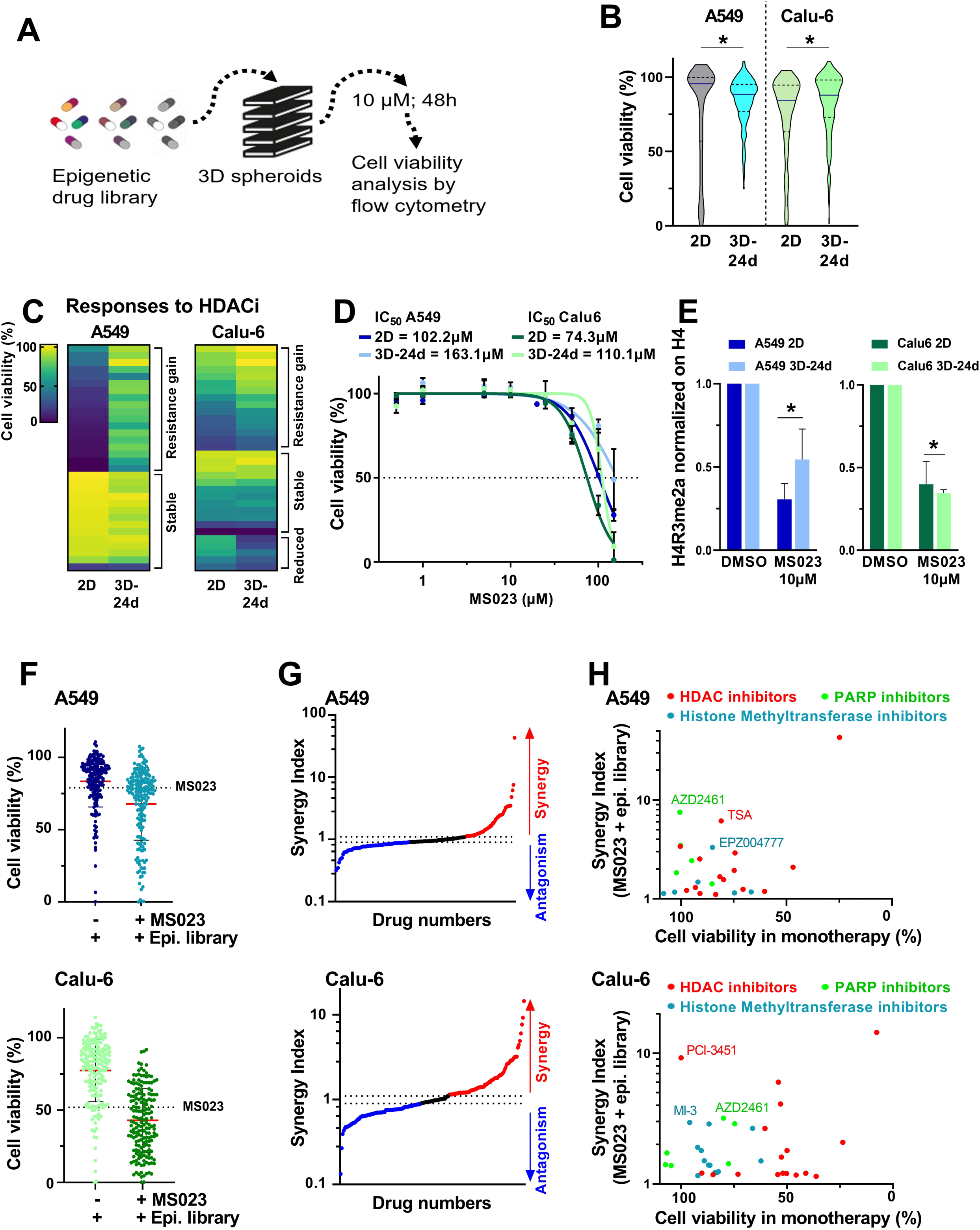
Long-term 3D spheroids reveal epigenetic vulnerability to type I PRMT inhibition in NSCLC. **a**) Drug screening of an epigenetic library composed of 181 compounds (10 µM, 48 hours treatment), in A549 and Calu-6 cells in 2D and 3D spheroids after 24 days (3D-24d). **b**) Violin plots of cell viability distribution after treatment. **c**) Heatmaps of cell viability distribution in response to 32 histone deacetylase (HDAC) inhibitors (10 µM, 48 hours treatment). **d**) Dose-response curves of type I PRMT inhibitor, MS023 in A549 and Calu-6 cells in 2D and 3D spheroids after 24 days. IC_50_ values are indicated. **e**) Western blot quantification of H4R3me2a levels in A549 and Calu-6 cells, in 2D and 3D-24d after MS023 treatment of 10 µM for 48 hours. Expression was normalized on H4 levels. **f**) Cell viability distribution of A549 3D-d24 (Top panel) or Calu-6 3D-d24 (Bottom panel), after 48 hours of epigenetic library compound in combination with MS023 (100 µM). Dotted line indicated the MS023 effect alone. **g**) Synergistic and antagonistic drug interactions between M023 and 181 epigenetic compounds. Top panel shows the results of A549 3D-24d and bottom panel shows the results on Calu-6 3D-24d. **h**) Synergy index of compounds from the most synergistic drug classes. Data are expressed relative to the cell viability of 3D-24d in monotherapy. Top panel shows the results of A549 3D-24d and bottom panel shows the results on Calu-6 3D-24d.

Due to the significant increase of H4R3me2a level in all long-term spheroid models (in A549, Calu-6 and Calu-1 cells), and its relevance in NSCLC, we thought of modulating its levels by inhibiting PRMT1, the type I protein methyltransferase, which catalyzes the reaction. We used the small-molecule compound MS023^61^ and determined drug sensitivity of A549 and Calu-6 cells in 2D and long-term spheroids (at day 24) after 48 hours treatment. Spheroids were more resistant to 2D cells (IC_50_: A549-2D = 102 µM; A549-3D-24d = 163 µM; Calu-6-2D = 74 µM; Calu-6-3D-24d = 110 µM; Figure 6d). Interestingly, even a non-toxic dose of MS023 (10 µM, 24 hours) was sufficient to reduce by 50% H4R3me2a level (measured by Western blotting) in both cell lines (Figure 6e and Figure S10a, Supporting information). Then, we reasoned that reducing H4R3me2a level through PRMT1 inhibition in long-term spheroids may prime tumor cells to novel therapeutic vulnerabilities. Thus, we treated A549 and Calu-6 spheroids with MS023 (100 µM) in combination with the epigenetic library (10 µM) in combination for 48 hours (Figure 6f). We calculated the synergy index^61,75^ by normalizing cell viability of drug in combination with MS023 to cell viability of each drug alone (for the epigenetic library), and to MS023 treatment alone. Interestingly, we discovered 74 (41%) and 78 (43%) synergistic interactions, for A549 and Calu-6 spheroids, respectively. On the other hand, we found 50 (28%) and 74 (41%) antagonistic interactions (Figure 6g).

Among the most promising synergistic interactions with the type I PRMT inhibitor, we identified three classes of inhibitors in both models (Figure 6h and Figure S10b, Supporting information). In particular, MS023 was synergistic with PARP inhibitors in A549 and Calu-6 spheroids, supporting our previous study.^61^ In addition, histone methyltransferase inhibitors were synergistic in combination with MS023 such as the G9a inhibitors (UNC0631 and BIX 01294); the Menin-MLL inhibitors (MI-2, MI-3), the SETD7 inhibitor (PFI-2) and the DOT1L inhibitors (SGC 0946 and EPZ004777). Most strikingly, we found that MS023 treatment synergized with HDAC inhibitors, suggesting that modulating arginine methylation levels may play a role in HDAC efficacy in the clinic. Overall, our protocol producing mature spheroids can be used for high-throughput drug screening, which lead to more stringent models than cells cultured in 2D for the drug discovery. Moreover, new synergistic drug interactions can be uncovered using long-term spheroids, which may bring novel therapeutic strategies for designing new clinical trials.

### Transcriptomic and chromatin changes gained during 3D culture were reversed by cultivating spheroids back to 2D culture

Lastly, we asked whether the transcriptomic and chromatin features acquired during long-term spheroid culture, which resemble more NSCLC tumors *in situ*, are stable or could be rapidly lost upon changes of cell culture method. To answer this question, we designed a protocol whereby NSCLC cells (A549, Calu-6 and Calu-1) from 3D spheroids at day 24 were dissociated by trypsinization and cultivated back in 2D for 7 days (referred to as 3D to 2D condition). After a week of 2D culture, we performed scRNA-seq, epigenetic analyses, and drug screening (Figure 7a). First, we performed scRNA-seq on A549 cells from the 3D to 2D conditions and compared the results to the A549 cells in 2D, and in spheroids after 3, 10, and 24 days. Surprisingly, only after 7 days in 2D, the 3D to 2D cells clustered close to the cells in 2D and opposite to the 24 days in 3D spheroids, suggesting a rapid return to the transcriptomic profile of 2D cells (Figure 7b). Importantly, this result strengthens the importance of cell culture conditions regarding transcriptomic profiles. Using SCENIC (Single-Cell regulatory Network Inference and Clustering) software, we identified a series of differentially expressed regulons between the different culture conditions of A549 cells (Figure 7c). Similar to bulk RNA-seq and scRNA-seq, the transition from 2D to 3D created a rapid shift in expression levels, which was the case for those transcription factors. Strikingly, the 3D to 2D conditions induced, in one only week, the reversal of expression profile that was acquired during 3 weeks of 3D culture, further supporting the impact of culture conditions on gene expression.

**Figure 7:**
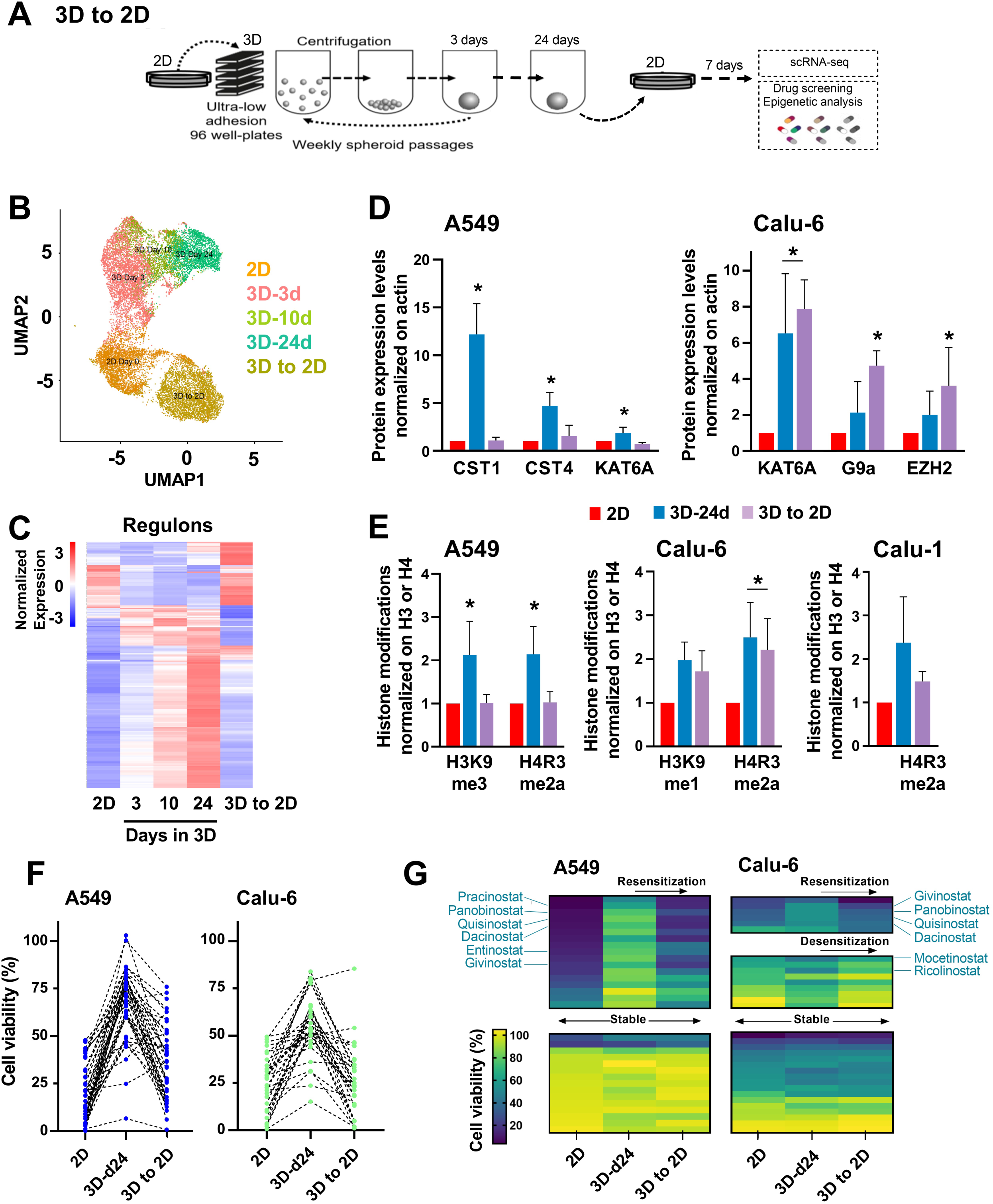
Transcriptomic signatures, chromatin modifications and pharmacological responses acquired during 3D culture were reversed by returning to 2D culture. **a**) Schematic representation of the experimental design to grow back 3D spheroids to 2D culture conditions. **b**) UMAP of A549 cells scRNA-seq data in 2D, 3D after 3, 10, 24 days and 3D to 2D. 3D to 2D refers to cells growth for 24 days in 3D and then plated in 2D petri dishes for 7 days. **c**) Heatmap of the most differentially expressed transcription factors in A549 2D, 3D after 3, 10, 24 days and 3D to 2D. **d**) Quantification of protein expression levels from western blots, in 3D spheroids (24 days) and 3D to 2D, relative to 2D condition in A549 (left panel) and Calu-6 (right panel) (mean ± SEM, n = 3; *p* < 0.05 by ANOVA). Actin was used as control. **e**) Quantification of post-translational modifications on histone H3 and H4 proteins of A549 (left panel), Calu-6 (middle panel) and Calu-1 (right panel) in 3D-24d and 3D to 2D relative to 2D (mean ± SEM, n = 3; *p* <0.05 by ANOVA). Total H3 and H4 were used as loading controls. **f**) Cell viability distribution after treatment with epigenetic drug library (10 µM, 48 hours) on A549 and Calu-6 in 2D, 3D-24d and 3D to 2D. Only the drugs inducing more that 50% death in 2D conditions are show in the graph. Dotted line link the viability of each individual drug under the different culture conditions. **g**) Heatmaps of cell viability distribution in response to HDAC inhibitor treatments (10 µM for 48 hours).

To evaluate whether the 3D to 2D conditions will impact not only transcriptomic profiles but also protein levels and chromatin modifications. We measured, by Western blotting, expression levels of CST1, CST4 and KAT6A in A549 spheroids, which gained expression in spheroids over time (Figure 2b and Figure 4e). Interestingly, protein expression levels returned to 2D levels in the 3D to 2D condition (Figure 7d). Similarly, histone modifications (H3K9me3 and H4R3me2a) followed this reversal when A549 cells were cultivated in 3D to 2D. The trend was observed for H4R3me2a in Calu-1 cells (Figure 7e). However, the proteins and histone marks that we investigated in Calu-6 cells were not reversed to 2D levels in the 3D to 2D condition, suggesting that either altered kinetics may occur with different cell models or other proteins could be impacted (Figure 7d-e).

Finally, in addition to transcriptomic and epigenetic changes, we asked whether the 3D to 2D culture condition could induce different pharmacological responses. To do so, we exposed A549 and Calu-6 cells, in 2D, in spheroids (after 24 days), and in the 3D to 2D condition to the epigenetic library (181 compounds) during 48 hours at 10 µM. Interestingly, both NSCLC cell lines showed sensitivity in 2D and resistance in spheroids, while in 3D to 2D conditions they showed the same sensitivity as in 2D (Figure 7f). Focusing on HDAC inhibitors, spheroids (of both cell lines) gained resistance to HDAC inhibitors, while 3D to 2D conditions resulted in resensitization to these inhibitors. Overall, gene expression changes and drug response acquired over 24 days in 3D culture were reversed after only 7 days of culture in 2D. Collectively, these data support that cell culture conditions play a crucial role on cell behavior, supporting well-characterized 3D spheroid modeling and well-established methods to ensure adequate preclinical studies.

## DISCUSSION

Cancer cell spheroids have been used for decades with the assumption that 3D culture may represent a more representative preclinical approach than classical 2D culture because of its 3D architecture, similar to human tumors. However, there are multiple approaches to perform 3D culture using either single or multiple spheroids per well, with or without extracellular matrix (like Matrigel or collagen) or growth factors, or dissociation steps (using trypsin or other methods). Thus, the lack of standardized or well-characterized methods to grow 3D cancer spheroids may render 3D culture more or less representative of human tumors regarding their transcriptomic and epigenomic landscapes and consequently to the pharmacological responses. Here, we used 96-well ultra low-adhesion plates, which are coated to prevent cell attachment to the plastic.^76^ These plates facilitate cell-cell interactions, and contribute to the formation of a spheroid structure.^77^ In addition, this technical approach is also adaptable to high-throughput drug screening.^12^

In the present study, we generated 3D cultures of NSCLC cell lines and we demonstrated that 3D cancer spheroids undergo a time-dependent transcriptomic, epigenomic, and pharmacological maturation process. After 24 days of 3D culture, we demonstrated that cancer cells adapted to the 3D culture, and better mimicked tumors *in situ*. This time-dependent transcriptomic and epigenomic reprogramming mirrored both transcriptomic pathways and epigenetic features that are playing key roles in NSCLC tumors. In particular, chromatin modifications were deeply affected both at methylation (on lysine and arginine residues) and acetylation levels. Other chromatin marks could be also impacted but will be to be explored in more details.

Generally, 3D spheroids are considered more resistant than classical 2D culture. Thus, we asked if transcriptomic and epigenetic changes occurring in 3D would translate into different pharmacological profiles over time. We chose to use a drug library of epigenetic/anticancer compounds since differential response between 2D and 3D spheroids could showcase not only changes in drug sensitivity but also could highlight epigenetic changes induced by 3D culture. This difference in sensitivity was illustrated by the resistance acquired to HDAC inhibitors in our NSCLC spheroid models. Interestingly, the HDAC inhibitors family is an epigenetic drug class, which demonstrated promising anticancer results in many preclinical studies, which lead to multiple clinical trials.^78^ Although few HDAC inhibitors are approved for hematological disorders, their preclinical efficacy against NSCLC preclinical models did not translate in clinical trials, since as single-agent therapy, only few responses and toxicity were observed.^79^ Interestingly, our long-term 3D NSCLC models would have predicted the failure of HDAC inhibitors in the clinic.^80^ However, we revealed that decreasing H4R3me2a in spheroids with MS023, a type I PRMT inhibitor, could sensitize NSCLC to HDAC inhibitors and other drug classes including PARP^61,81^ and some histone methyltransferase inhibitors, providing the rationale to start novel clinical investigations. Indeed, PRMTs and histone methyltransferases have been shown to interact together.^82^ However, our data show that such interaction is more critical to NSCLC survival in mature spheroids.

Lastly, we demonstrated by cultivating the 3D spheroids back into 2D conditions that the transcriptomic and epigenomic and pharmacological responses gained after 24 days on 3D culture could be lost only after one week back into 2D culture, suggesting the detrimental impact of classical 2D culture. These results highlight the importance of tumor microenvironment to cancer cell phenotype and how is also influence drug response.^9^ Although many studies demonstrated the gain of tumor features in 3D culture, our results show how rapidly those gains can be lost upon cancer cell growth back in 2D.

One of the main limitations in development of anticancer drugs is the lack of correlation of preclinical results generally obtained from 2D cell cultures and xenograft models from bench to bedside.^6,83^ One of the most relevant and advantage of cancer spheroids is drug screening, as they allow fast, robust and affordable drug screening models. In line with our findings, using well-characterized 3D spheroids (supported with transcriptomic/epigenomic datasets and their correlation to patient cohorts) may help developing more predictive *in vitro* cancer models for the drug discovery pipeline, which may increase the translational success of preclinical studies. Our work is in line with the current use of well-annotated and characterized patient-derived organoids.^84–87^

Despite the interesting properties of spheroids, several challenges still hamper their broader use as preclinical tools for therapeutic efficacy evaluation for new anticancer agents. One of the main issue relates to the absence of standardized culture methods. Here, we propose a 3D culture method allowing the generation of spheroids, which are more predictive models in NSCLC. The recent passage of the FDA Modernization Act 2.0 (in 2022), now allows the replacement of certain animal studies by using alternative models such as spheroids, which could cause a surge in the popularity of these 3D spheroid models. This is particularly important as we demonstrated that transcriptomic and epigenomic profiles in spheroids could be easily reversed by changing conditions, suggesting that, at least, some of these features may not be maintained in animal models.

In summary, we demonstrated that 3D cancer spheroids undergo a time-dependent transcriptomic, epigenomic, and pharmacological maturation process, which better mimic NSCLC tumors *in situ*. This maturation process observed during 3D culture could be exploited in other solid tumor preclinical models to improve *in vitro* preclinical basic and pharmacological studies. This maturation process observed during 3D culture could be exploited in other solid tumor preclinical models to improve *in vitro* preclinical basic and pharmacological studies. Drug discovery in oncology is a long and risky process, which involves high-throughput drug screenings, hit identification, preclinical validations, and finally clinical trials. Despite tremendous investments, exceeding USD 1 billion Dollars and more than a decade of efforts in R&D, the probability of a given anticancer drug that enter phase I clinical trial to obtain regulatory approval is still around 5%.^6,83^ Among the most common factors responsible for drug failure during clinical development are the lack of clinical efficacy, which occurred in more than half of the studies (57%), followed by commercial reasons (22%) and toxicity issues (17%).^88^ Altogether, the high failure rate in drug development highlights the complexity to develop new drugs in oncology and more strikingly, that achieving therapeutic prediction from preclinical research to clinical trials is a difficult endeavour.^6,83^ Thus, using well-characterized 3D spheroids that undergo a time-dependent transcriptomic and epigenomic maturation process mediated by the 3D microenvironment may represent robust models that faithfully recapitulate the human tumors, allowing target identification and predict for therapeutic efficacy in patients.

## DECLARATIONS

### Availability of data and material

All raw sequencing data files are available from the GEO database (www.ncbi.nlm.nih.gov/geo/); (currently in submission, accession numbers will be added before final submission)

### Competing interests

The Authors declare that they have no conflict of interest relevant the above manuscript.

### Funding

This work was funded by the Canadian Institute of Health Research (CIHR), the Natural Sciences and Engineering Research Council of Canada (NSERC) and Foundation Charles-Bruneau. Single-cell sequencing costs were funded by Illumina-CHU Sainte-Justine Research centre. Full Reference Epigenomic sequencing of A549 cell line grown in 2D and 3 time points in 3D was funded by Community Access Program from the Canadian Epigenetics Environment and Heath Research Consortium (CEEHRC). N. J.-M. R. and S.McG hold research scholar senior awards from the Fonds de Recherche du Québec en Santé.

### Authors’ contributions

A. D., N. S., G. McI., and S. L. performed *in vitro* experiments, analyzed results and wrote the manuscript. M. H., A. M., A. L., O. V., F. M., M. C., P. S-O, and G. C. performed bioinformatics analyses and wrote the manuscript. G. A., S. R., D. S., S. McG. analyzed results and edited the manuscript. N. J.-M. R. conceptualization, funding acquisition, methodology, project administration, resources, supervision, writing and review the manuscript.

## Acknowledgements

The authors acknowledge Dr. Claudia Kleinman from the Lady Davis Institute for her suggestions regarding the single-cell sequencing analysis.

## SUPPLEMENTARY FIGURE LEGENDS

**Supplementary figure S1: Characterization of lung cancer 3D spheroids. a**) Left panel: A549 spheroid diameter (µm) at different initial seeding densities (5000, 20,000, 40,000 cells per well). Right panel: Calu-1 and Calu-6 spheroid area. Spheroids were seeded with 20,000 cells and 10,000 cells per well, respectively. Diameters and areas were measured every 6 or 8 hours for the first 2 days and then after 24 hours with the Live Cell Imaging System Incucyte™ (mean ± SD, n = 3). **b**) Representative bright field microscopy images of Calu-1 cells in 3D spheroid culture at day 0 (immediately after spheroid assembly), day 1, day 3 and day 10. Pictures were taken with Live Cell Imaging System Incucyte™. Scale bar for all images represents 800 µm. **c**) Frequency distribution of 96 A549 spheroid areas (µm2) after 0, 1, 3 and 7 days in 3D. Spheroid area was measured by Live Cell Imaging System Incucyte™. **d**) Number of A549 cells per spheroid for the first 5 days of 3D culture with different initial seeding densities (5000, 20,000, 40,000 cells per well). Cells were trypsinized and then counted manually by haemocytometer with trypan blue staining (mean ± SD, n = 3). **e)** Doubling times (hours) of A549 cells in 2D and 3D spheroid culture after 3, 10, 17, 24, 31, 38 days. **f**) Cell viability of Calu-1 2D and spheroids after 1, 3, 10, 17, 24 days in 3D culture using cell viability staining (ViaCount™ reagent) as compared to 2D condition. Statistically significant changes are indicated by a star (p < 0.05 by paired ANOVA).

**Supplementary figure S2: Time-dependent changes in gene expression changes during 3D culture, by bulk RNA-seq. a**) Heatmap showing expression levels of 10,000 most differentially expressed genes obtained by bulk RNA-sequencing in A549 cells in 2D culture and after 3, 10 and 24 days in spheroids (n = 3). **b**) Principal component analysis of A549 cells in 2D culture and after 3, 10 and 24 days in spheroids (n = 3). **c**) Gene expression relative changes from 2D culture to 3D culture at the different time points. A linear regression was calculated to estimate gene expression changes in the different time points. Positive and negative estimate values correspond to an increase or decrease expression from 2D to 3D, respectively. **d**) Gene set enrichment analysis using C6 oncogenic gene sets. **e**) Gene set enrichment analysis using C5 molecular function and C5 biological process gene sets.

**Supplementary figure S3: Lung A549 spheroids gene expression changes and EMT phenotype. a-d)** Heatmaps representing gene expression changes during 3D culture related to **a**) hypoxia, **b**) KRAS, **c**) G2/M and **d**) epithelial-mesenchymal transition pathways (EMT). RNA-seq results were obtained from A549 cells in 2D and in 3D after 3, 10 and 24 days in culture. **e-f)** Gene expression analysis by qPCR of EMT markers: **e**) N-cadherin and E-cadherin; **f**) Snail, and Slug. RNA was harvested from A549 cells in 2D and after 3, 10, 17, and 24 days in 3D. Expression is relative to the levels in 2D cells (mean ± SEM, n = 3). **g**) Transwell invasion assay of A549 cells in 2D and after 10 and 24 days in 3D. Images were taken under bright field DMi6 Leica microscope (5X objective). Invasion index was calculated from five distinct fields of same image (mean ± SEM, n = 3; a star indicates *p* < 0.05; One-way Anova).

**Supplementary figure S4: Epigenetic changes during 3D spheroid culture of A549 cells. a**) Protein expression levels of HDAC1, HDAC2, HDAC3, HDAC4, HDAC5, KAT2B and KAT5 by western blotting. Actin was used as a loading control. Histograms show quantification of protein expression relative to 2D control (mean ± SEM, n = 3; a star indicates *p* <0.05; Two-way Anova). **b**) Representative western blots images showcasing levels of H3K9ac, H4K16ac, H3K9me1, H3K9me2, H3K9me3, H3K27me3 in 2D and after 3, 10, 17, and 24 days in 3D. Total H3 and H4 were used as loading controls. **c-d**) Quantification of acetylation levels on H3K14, H3K18, H3K27, H3K56, H4K5, H4K8 and H4K12 by western blotting. Total H3 or H4 were used as loading controls (mean ± SEM, n = 3).

**Supplementary figure S5: Time-dependent changes in chromatin marks and impact of transcription factors expressed in 3D spheroids on survival of NSCLC patients.** Chromatin Immunoprecipitation of **a**) H3K4me1 and **b**) H3K36me3 marks centered on transcription start sites (TSS) in A549 cells in 2D and in spheroids after 3, 10 and 24 days. Probability of Overall Survival in function of the expression levels of **c**) CDH4, **d**) ELF3, **e**) ELF5, **f**) PU.1, and **g**) GATA1 using cBioPortal.

**Supplementary figure S6: Gene Set Enrichment analysis.** Gene sets of **a**) upregulated genes marked by H3K4me3 (n=799) in their promoter regions and **b**) upregulated genes marked by H3K27ac (n=533) in their promoter regions in A549 3D spheroids (after 24 days in culture).

**Supplementary figure S7: Whole genome bisulfite sequencing analyses of DNA methylation changes between 2D culture and 3D spheroids (at day 3, 10 and 24) in A549 cells.** Heatmaps show **a**) the intragenic DNA methylation levels and **b**) the promoter DNA methylation levels in A549 2D and 3D days 3, 10 and 24. **c**) Beta value expression changes from 2D to 3D (at day 24) in function of Beta value of DNA methylation. In each four quadrant, the number of gene is indicated.

**Supplementary figure S8: a**) Complete list of pathways from Figure 4D showing single-sample gene set enrichment analysis between the 15 different clusters. Adjusted *p*-values and normalized enrichment score scales are shown in the graph. **b**) Patient samples (n=18) proportion of normal vs malignant cells. **c**) Spearman correlation of malignant cells and normal cells from the 18 different patients with the A549 cells in culture (in 2D and 3D at the different time points).

**Supplementary figure S9: A549 and Calu-6 drug screening.** Cell viability distribution of **a**) A549 and **b**) Calu-6 in 2D and 3D-24d after treatment with 181 epigenetic drugs (treatment of 10µM, at 48h). **c**) Cell viability distribution clustered by target classes in A549 and Calu-6 in 2D and 3D-24d. Drug class names are indicated at the bottom of the figure.

**Supplementary figure S10: PRMT inhibition on NSCLC spheroids. a**) Representative western blotting images of H4R3me2a and H4 proteins in A549 and Calu-6 2D and 3D spheroids after 48h treatment with MS023 (1 µM, 10 µM or 20 µM). Calu-6 quantification for all doses are shown in the right panel. (n=3, *: *p* < 0.05). **B**) Cell viability distribution clustered by target classes of A549 3D-d24 (left panel) and Calu-6 3D-d24 (right panel), after 48 hours of epigenetic library compound in combination with MS023 (100 µM). Drug class names are indicated at the bottom of the figure.

**Supplementary Table S1:** Antibodies used for western blots.

| **Antibody** | **Dilution** | **Supplier / cat number** |
| --- | --- | --- |
| Actin-β | 1:5000 | Sigma-Aldrich cat#A2228 |
| BRG1 | 1 :1000 | Cell Signaling cat#49360 |
| Caveolin1 | 1 :1000 | Invitrogen cat#MA3-600 |
| CEACAM6 | 1 :1000 | Invitrogen cat#MA5-29144 |
| CST1 | 1 :1000 | Invitrogen cat#PA5-39123 |
| CST4 | 1 :1000 | Invitrogen cat#MA5-25828 |
| EZH2 | 1 :1000 | Invitrogen cat#MA5-18108 |
| G9a | 1 :1000 | Active Motif cat#91230 |
| H3 | 1:5000 | Active Motif cat#39763 |
| H3K14ac | 1:5000 | Active Motif cat#39698 |
| H3K18ac | 1:2500 | Active Motif cat#39588 |
| H3K27ac | 1:5000 | Active Motif cat#39134 |
| H3K27m3 | 1:2500 | Active Motif cat#39155 |
| H3K56ac | 1:5000 | Active Motif cat# 39282 |
| H3K9ac | 1:5000 | Active Motif cat#39917 |
| H3K9me1 | 1:5000 | Active Motif cat#39887 |
| H3K9me2 | 1:5000 | Active Motif cat#39753 |
| H3K9me3 | 1:5000 | Active Motif cat#39765 |
| H4 | 1:5000 | Active Motif cat#39270 |
| H4K12ac | 1:5000 | Active Motif cat#39928 |
| H4K16Ac | 1:5000 | Active Motif cat#39167 |
| H4K5ac | 1:2000 | Active Motif cat#39700 |
| H4K8ac | 1:5000 | Active Motif cat#39172 |
| H4R3me2a | 1:1000 | Active Motif cat# 39006 |
| HDAC1 | 1:2500 | Cell Signaling cat#5356 |
| HDAC2 | 1:1000 | Cell signaling cat#5113 |
| HDAC3 | 1:2000 | Cell Signaling cat#3949 |
| HDAC4 | 1:2500 | Cell Signaling cat#7628 |
| HDAC5 | 1:1000 | Active Motif cat#40970 |
| HDAC6 | 1:1000 | Cell Signaling cat#7558 |
| KAT2A | 1 :500 | Santa Cruz cat#sc-365321 |
| KAT3A | 1 :1000 | Cell Signaling cat#7389 |
| KAT3B | 1 :2000 | Active Motif cat#61402 |
| KAT5 | 1 :2500 | Abcam cat#ab137518 |
| KAT6A | 1 :1000 | Active Motif cat#39868 |
| KAT7 | 1 :1000 | Bethyl Labs cat#A302-224A-T |
| LSD1 | 1:1000 | Active Motif cat# 61608 |

**Supplementary Table S2:**
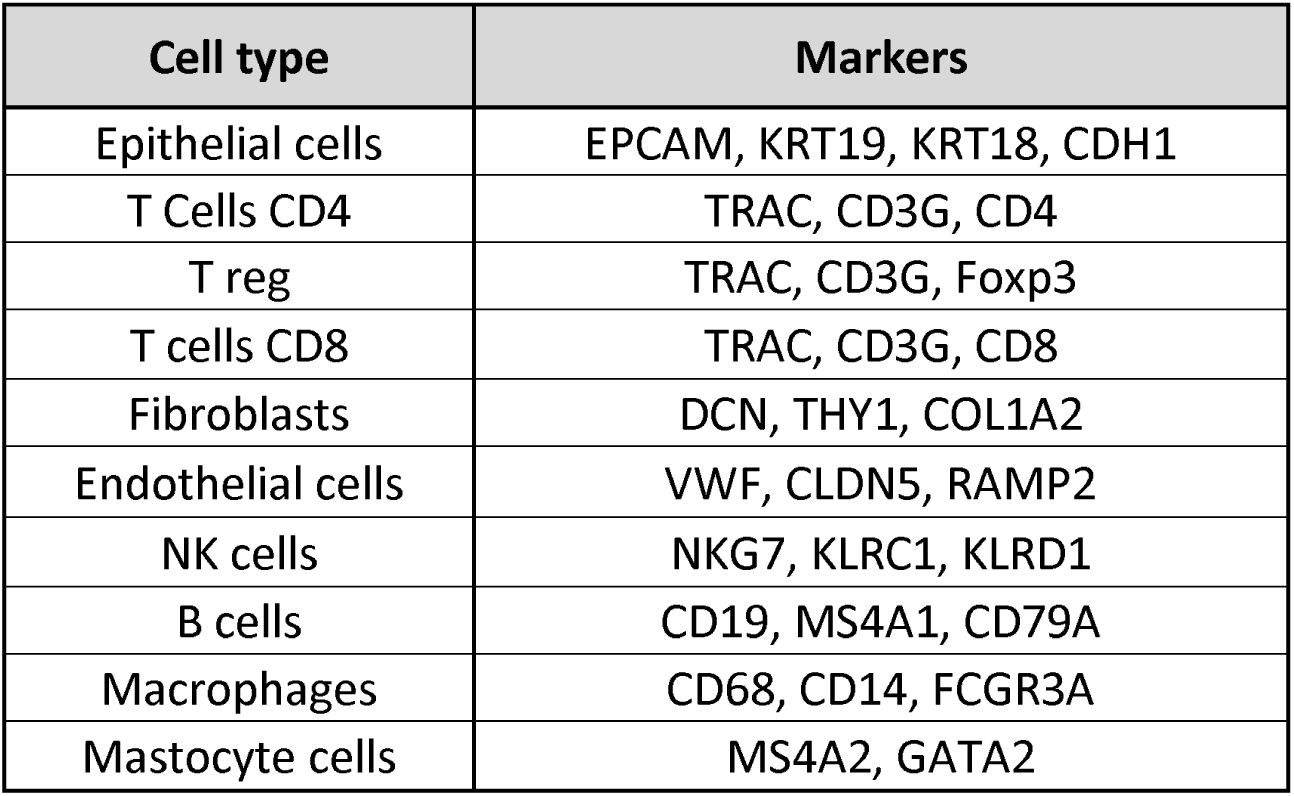
List of cell type specific markers used for sc-Seq analysis.

| Cell type | Markers |
| --- | --- |
| Epithelial cells | EPCAM, KRT19, KRT18, CDH1 |
| T Cells CD4 | TRAC, CD3G, CD4 |
| T reg | TRAC, CD3G, Foxp3 |
| T cells CD8 | TRAC, CD3G, CD8 |
| Fibroblasts | DCN, THY1, COL1A2 |
| Endothelial cells | VWF, CLDN5, RAMP2 |
| NK cells | NKG7, KLRC1, KLRD1 |
| B cells | CD19, MS4A1, CD79A |
| Macrophages | CD68, CD14, FCGR3A |
| Mastocyte cells | MS4A2, GATA2 |

**Supplementary Table S3:**
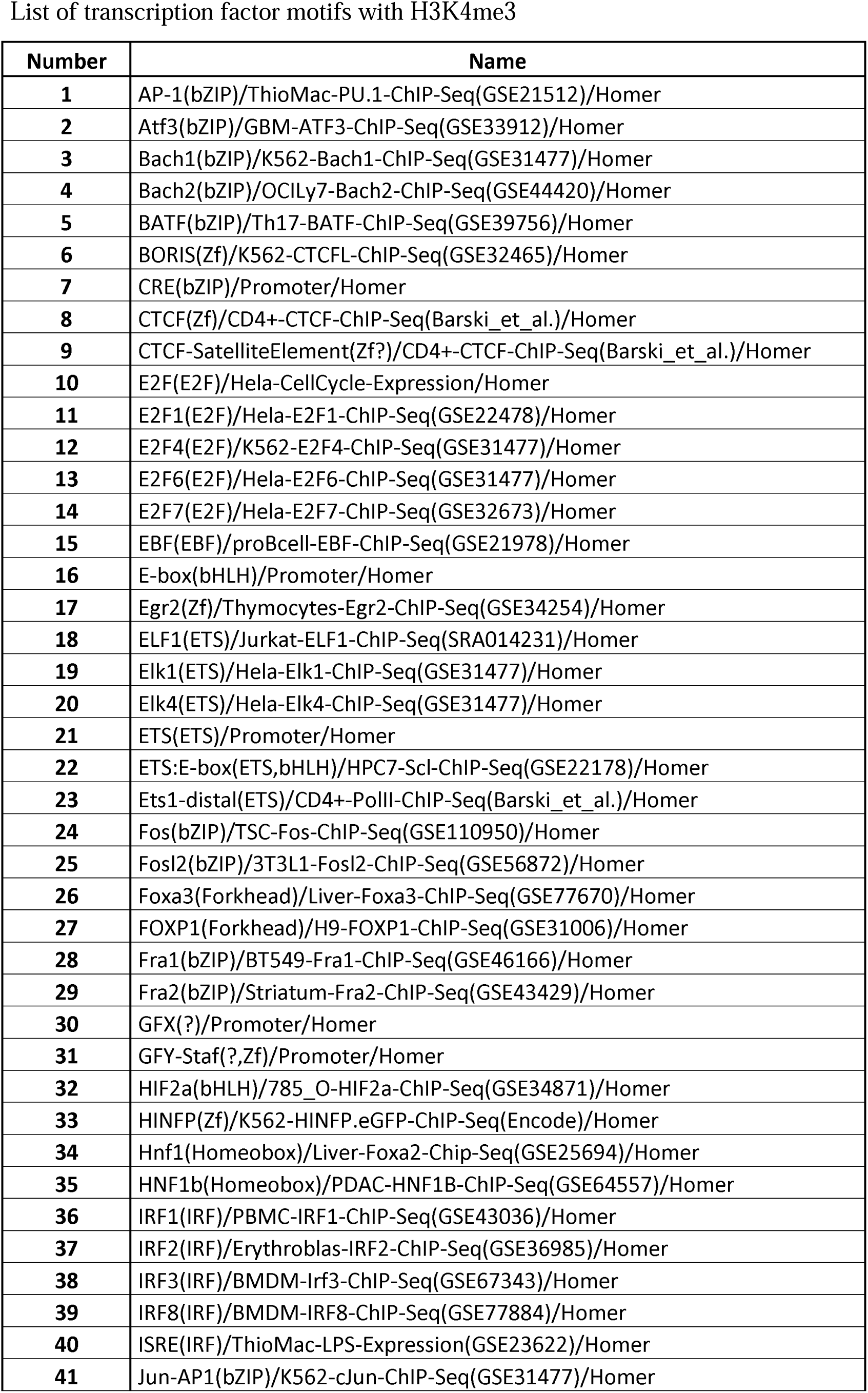
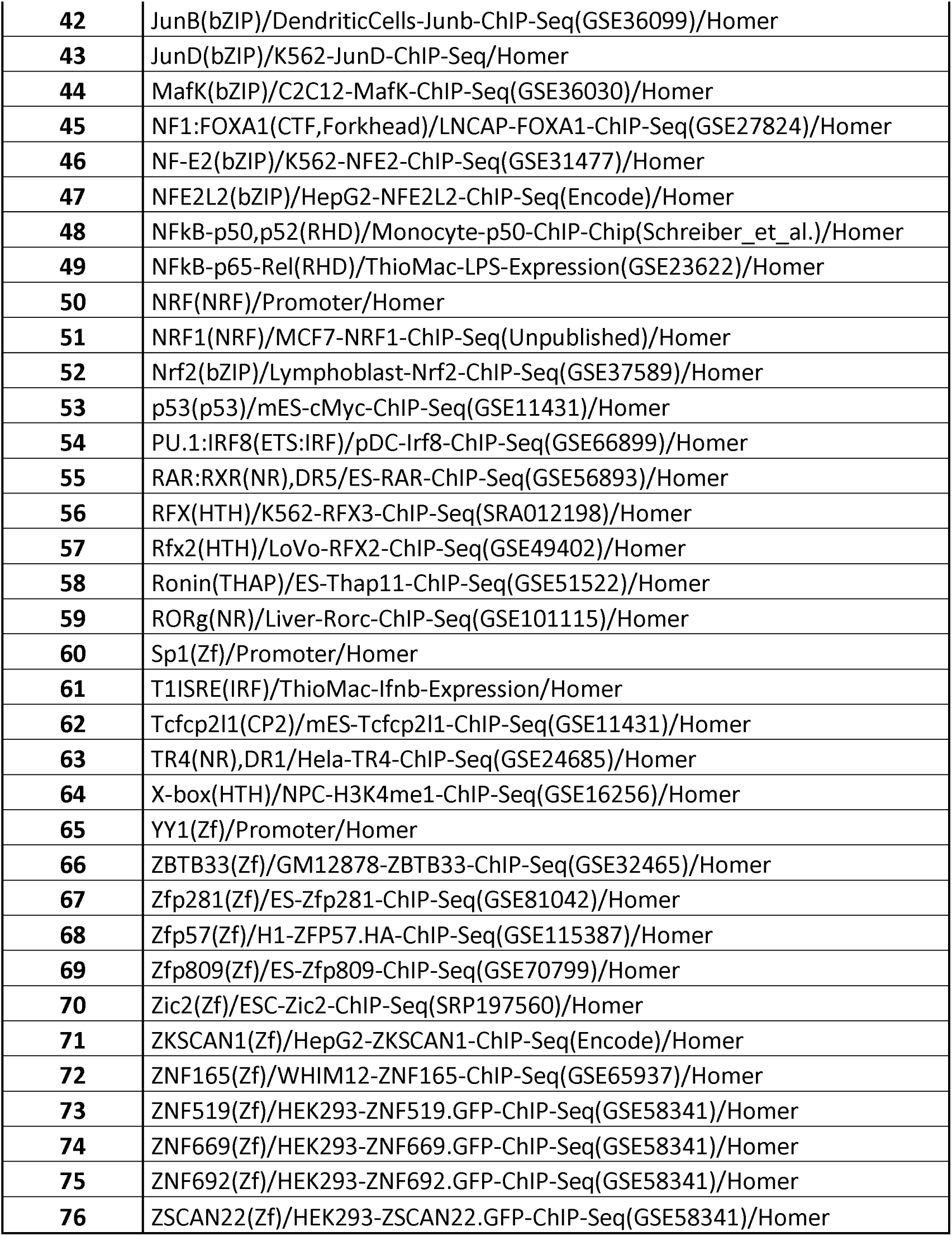
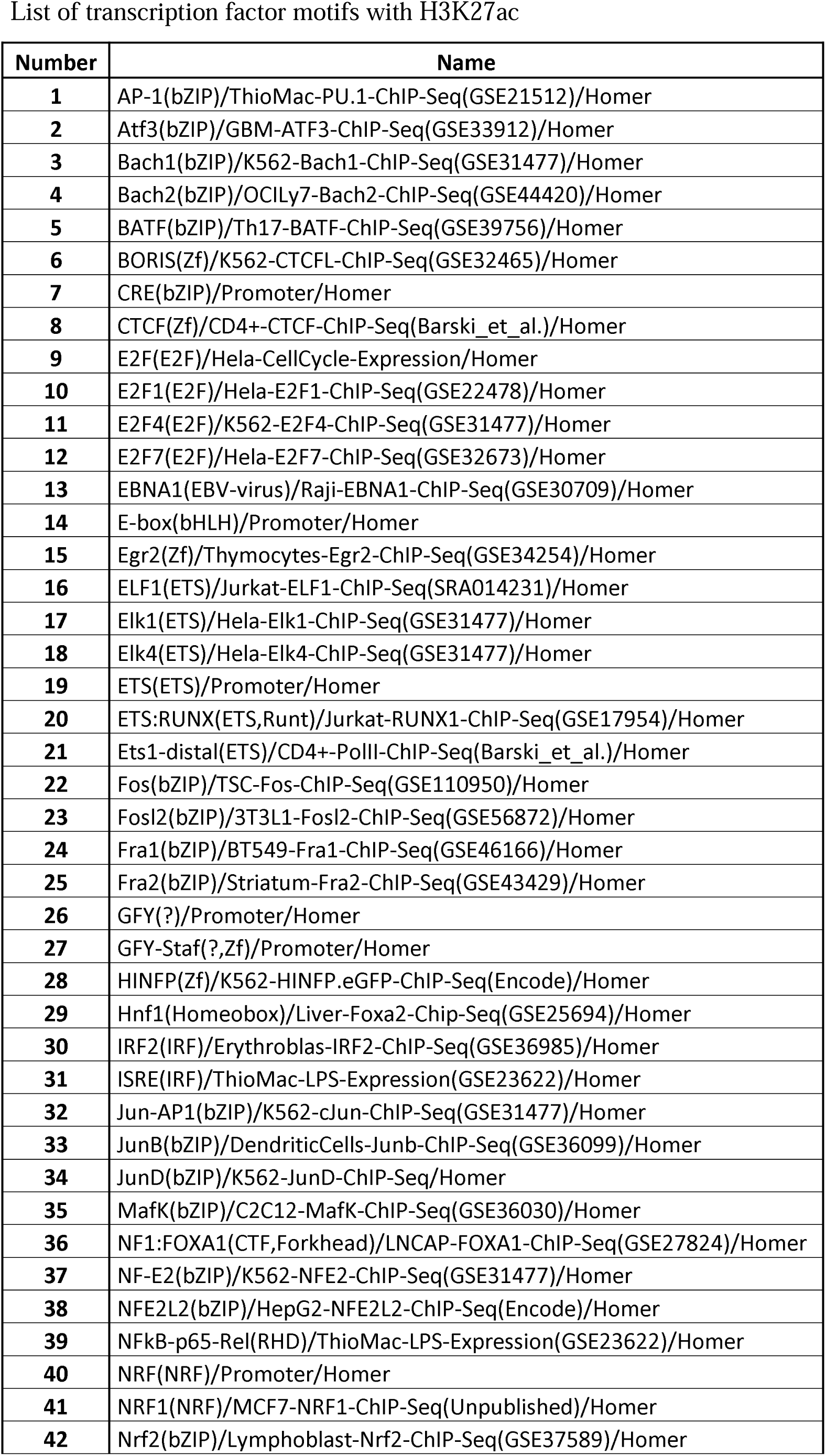
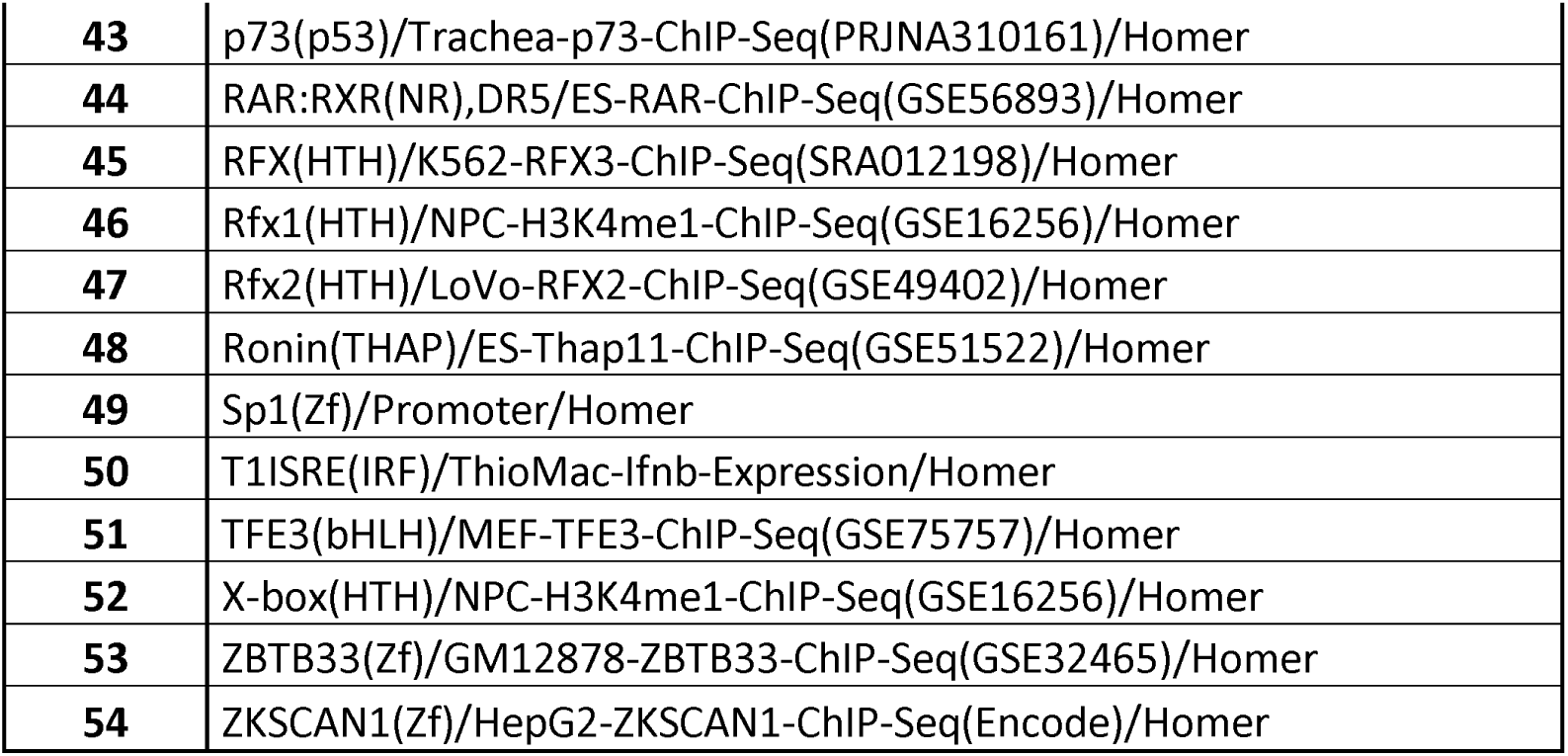
Lists of transcription factor motifs List of transcription factor motifs with H3K4me3.

## Notes

### Competing Interest Statement

The authors have declared no competing interest.

