## Supplemental Figures 1 to 10 for "Time-dependent chromatin maturation during 3D spheroid culture improves preclinical modeling of non-small cell lung cancer"

### Figure S1

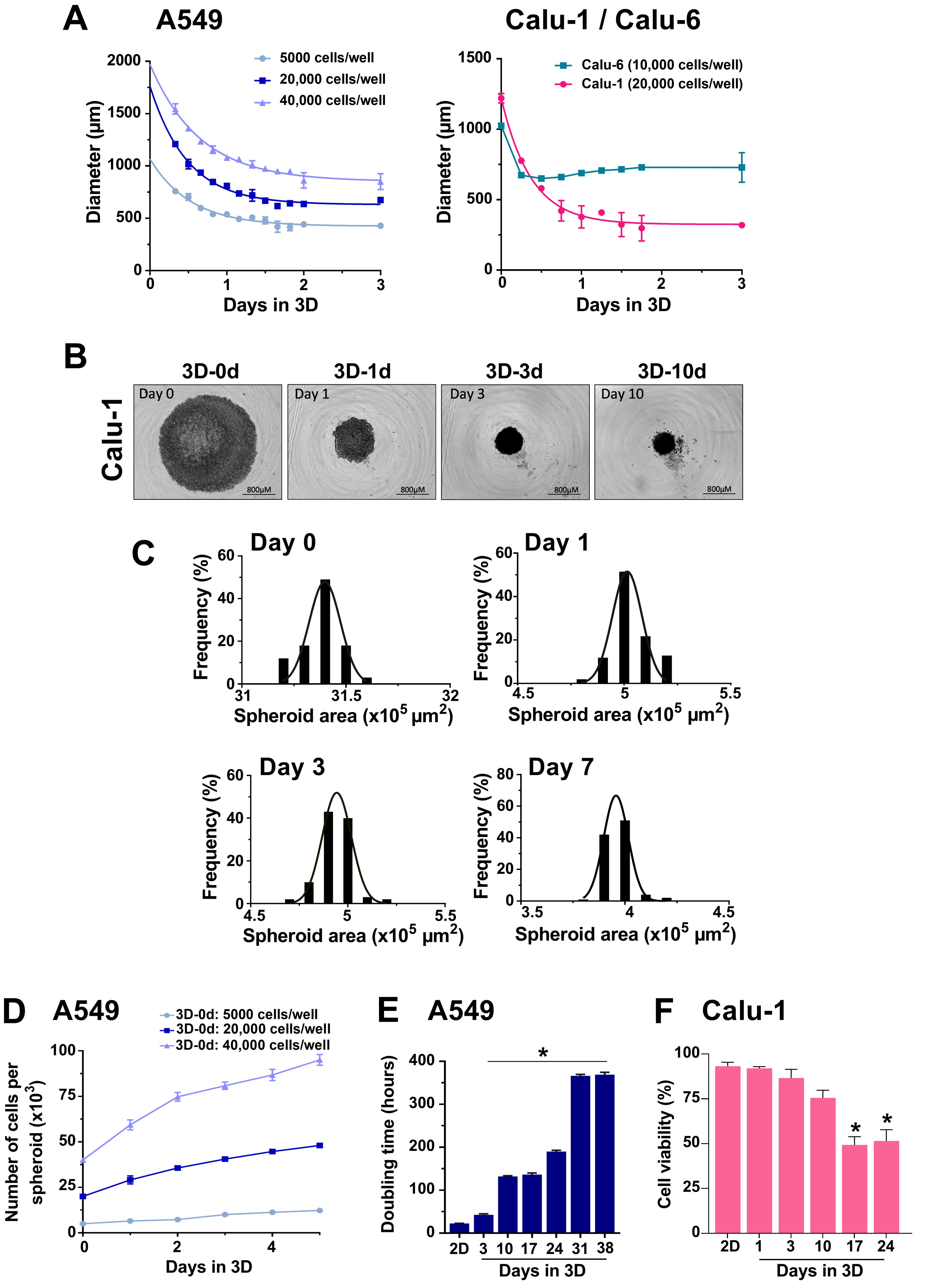

A
A549
Diameter (μm)
5000 cells/well
20,000 cells/well
40,000 cells/well
Days in 3D
Calu-1 / Calu-6
Calu-6 (10,000 cells/well)
Calu-1 (20,000 cells/well)
B
3D-0d
3D-1d
3D-3d
3D-10d
Calu-1
Day 0
Day 1
Day 3
Day 10
800μM
C
Day 0
Day 1
Day 3
Day 7
Frequency (%)
Spheroid area (x10^5 μm^2)
D
A549
3D-0d: 5000 cells/well
3D-0d: 20,000 cells/well
3D-0d: 40,000 cells/well
Number of cells per spheroid (x10^3)
Days in 3D
E
A549
Doubling time (hours)
2D
Days in 3D
F
Calu-1
Cell viability (%)
2D
Days in 3D

### Figure S2

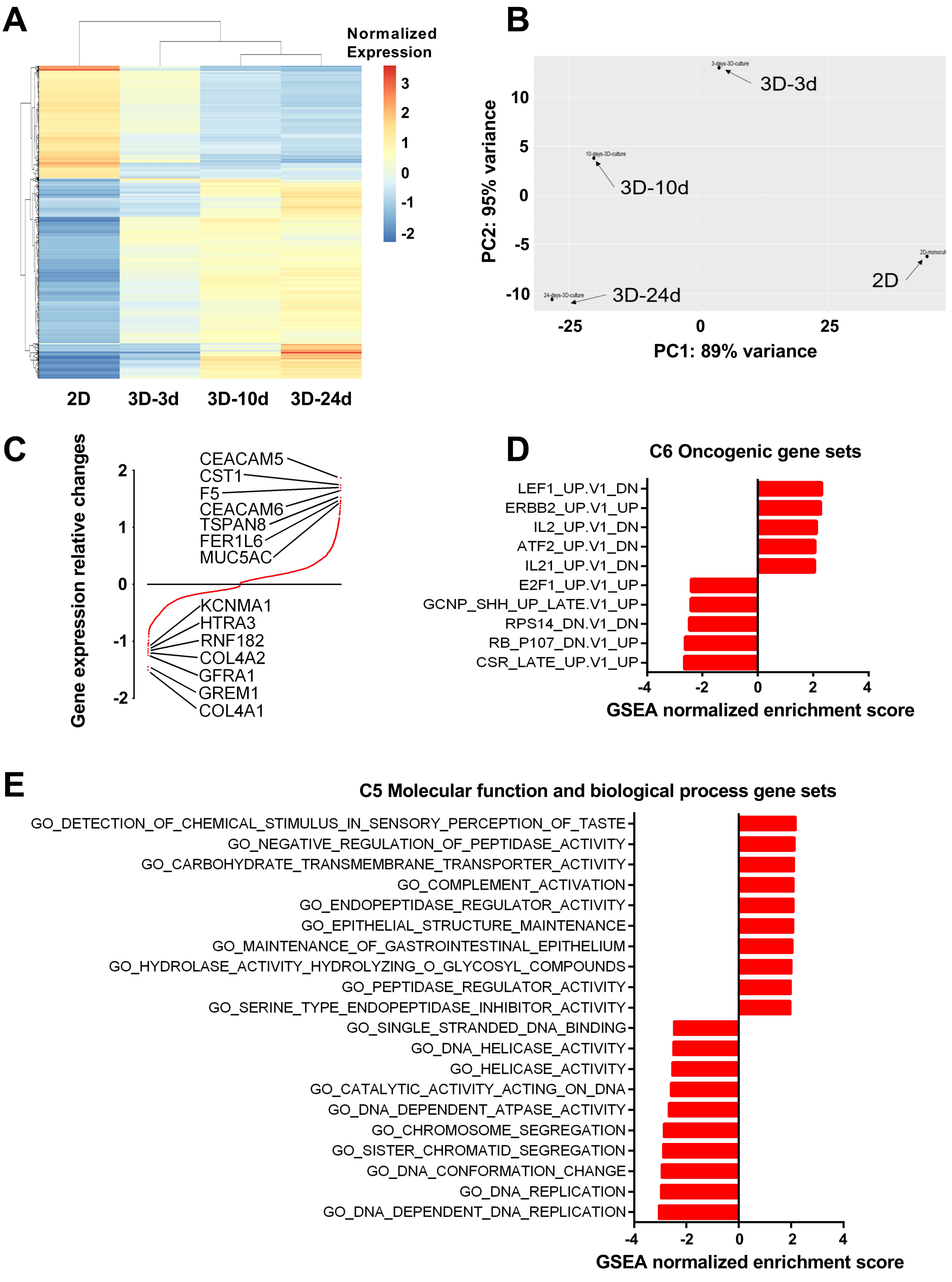

A
Normalized Expression
3
2
1
0
-1
-2
2D
3D-3d
3D-10d
3D-24d
B
3D-3d
3D-10d
3D-24d
2D
PC2: 95% variance
PC1: 89% variance
10
5
0
-5
-10
-25
0
25
C
Gene expression relative changes
CEACAM5
CST1
F5
CEACAM6
TSPAN8
FER1L6
MUC5AC
KCNMA1
HTRA3
RNF182
COL4A2
GFRA1
GREM1
COL4A1
2
1
0
-1
-2
D
C6 Oncogenic gene sets
LEF1_UP.V1_DN
ERBB2_UP.V1_UP
IL2_UP.V1_DN
ATF2_UP.V1_DN
IL21_UP.V1_DN
E2F1_UP.V1_UP
GCNP_SHH_UP_LATE.V1_UP
RPS14_DN.V1_DN
RB_P107_DN.V1_UP
CSR_LATE_UP.V1_UP
-4
-2
0
2
4
GSEA normalized enrichment score
E
C5 Molecular function and biological process gene sets
GO_DETECTION_OF_CHEMICAL_STIMULUS_IN_SENSORY_PERCEPTION_OF_TASTE
GO_NEGATIVE_REGULATION_OF_PEPTIDASE_ACTIVITY
GO_CARBOHYDRATE_TRANSMEMBRANE_TRANSPORTER_ACTIVITY
GO_COMPLEMENT_ACTIVATION
GO_ENDOPEPTIDASE_REGULATOR_ACTIVITY
GO_EPITHELIAL_STRUCTURE_MAINTENANCE
GO_MAINTENANCE_OF_GASTROINTESTINAL_EPITHELIUM
GO_HYDROLASE_ACTIVITY_HYDROLYZING_O_GLYCOSYL_COMPOUNDS
GO_PEPTIDASE_REGULATOR_ACTIVITY
GO_SERINE_TYPE_ENDOPEPTIDASE_INHIBITOR_ACTIVITY
GO_SINGLE_STRANDED_DNA_BINDING
GO_DNA_HELICASE_ACTIVITY
GO_HELICASE_ACTIVITY
GO_CATALYTIC_ACTIVITY_ACTING_ON_DNA
GO_DNA_DEPENDENT_ATPASE_ACTIVITY
GO_CHROMOSOME_SEGREGATION
GO_SISTER_CHROMATID_SEGREGATION
GO_DNA_CONFORMATION_CHANGE
GO_DNA_REPLICATION
GO_DNA_DEPENDENT_DNA_REPLICATION
-4
-2
0
2
4
GSEA normalized enrichment score

### Figure S3

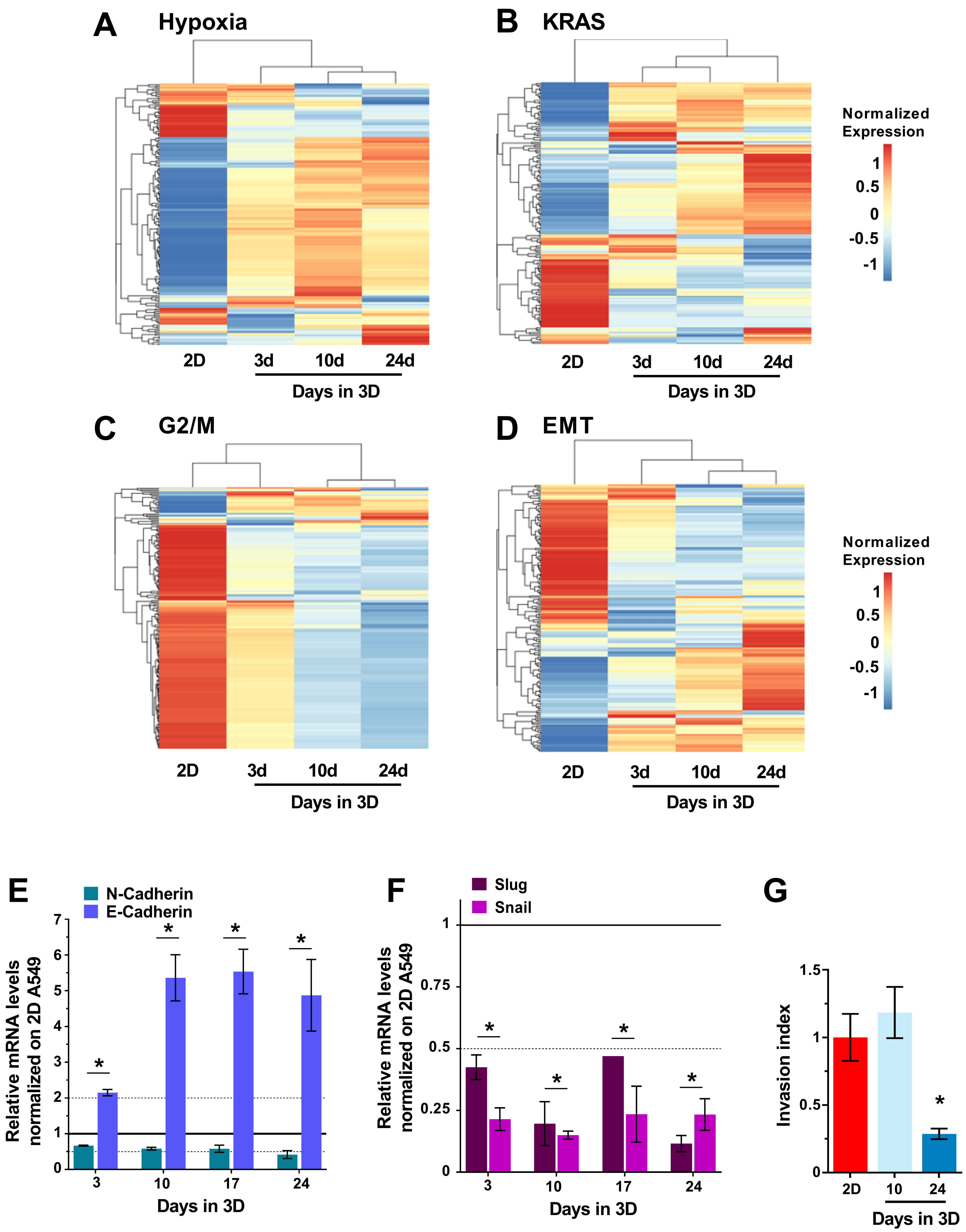
A
Hypoxia
2D
3d
10d
24d
Days in 3D
B
KRAS
2D
3d
10d
24d
Days in 3D
Normalized
Expression
1
0.5
0
-0.5
-1
C
G2/M
2D
3d
10d
24d
Days in 3D
D
EMT
2D
3d
10d
24d
Days in 3D
Normalized
Expression
1
0.5
0
-0.5
-1
E
N-Cadherin
E-Cadherin
Relative mRNA levels
normalized on 2D A549
7
6
5
4
3
2
1
0
3
10
17
24
Days in 3D
F
Slug
Snail
Relative mRNA levels
normalized on 2D A549
1
0.75
0.5
0.25
0
3
10
17
24
Days in 3D
G
Invasion index
1.5
1
0.5
0
2D
10
24
Days in 3D

### Figure S4

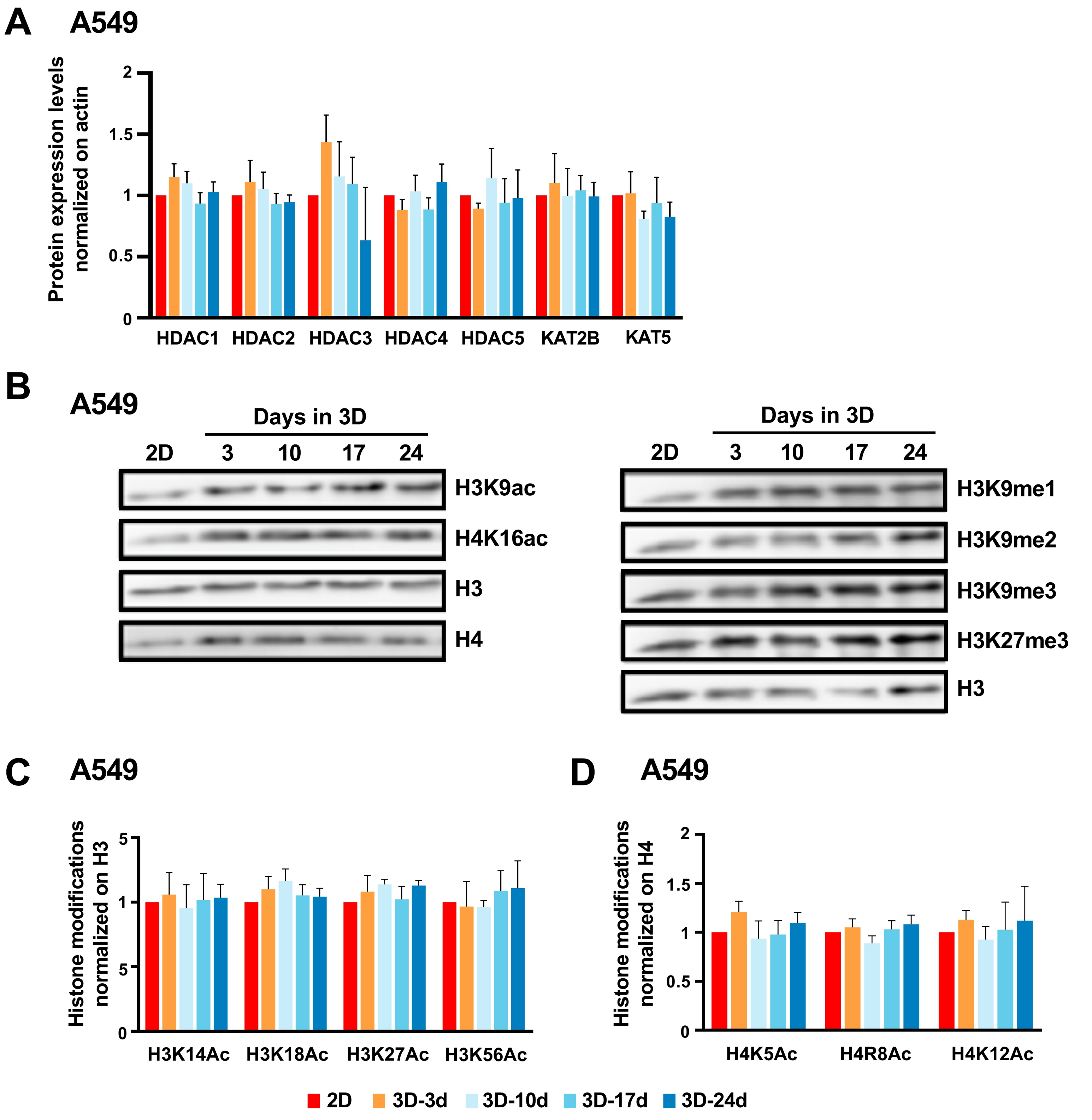
A
A549
Protein expression levels normalized on actin
HDAC1
HDAC2
HDAC3
HDAC4
HDAC5
KAT2B
KAT5
B
A549
Days in 3D
2D
3
10
17
24
H3K9ac
H4K16ac
H3
H4
H3K9me1
H3K9me2
H3K9me3
H3K27me3
C
A549
Histone modifications normalized on H3
H3K14Ac
H3K18Ac
H3K27Ac
H3K56Ac
D
A549
Histone modifications normalized on H4
H4K5Ac
H4R8Ac
H4K12Ac
2D
3D-3d
3D-10d
3D-17d
3D-24d

#### Figure S5

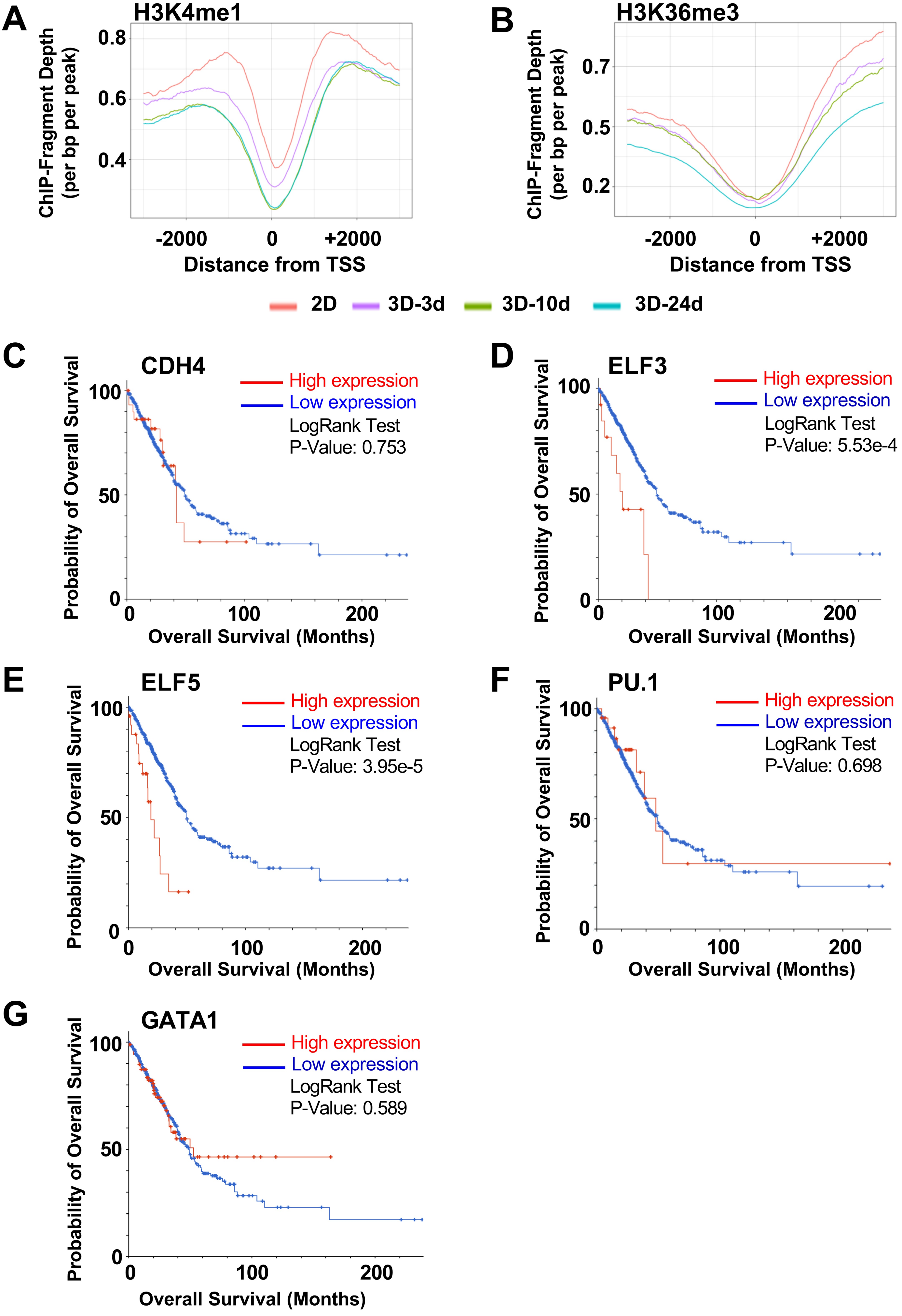
A
H3K4me1
ChIP-Fragment Depth (per bp per peak)
0.8
0.6
0.4
-2000
0
+2000
Distance from TSS
B
H3K36me3
ChIP-Fragment Depth (per bp per peak)
0.7
0.5
0.2
-2000
0
+2000
Distance from TSS
2D
3D-3d
3D-10d
3D-24d
C
CDH4
High expression
Low expression
LogRank Test
P-Value: 0.753
Probability of Overall Survival
100
50
0
0
100
200
Overall Survival (Months)
D
ELF3
High expression
Low expression
LogRank Test
P-Value: 5.53e-4
Probability of Overall Survival
100
50
0
0
100
200
Overall Survival (Months)
E
ELF5
High expression
Low expression
LogRank Test
P-Value: 3.95e-5
Probability of Overall Survival
100
50
0
0
100
200
Overall Survival (Months)
F
PU.1
High expression
Low expression
LogRank Test
P-Value: 0.698
Probability of Overall Survival
100
50
0
0
100
200
Overall Survival (Months)
G
GATA1
High expression
Low expression
LogRank Test
P-Value: 0.589
Probability of Overall Survival
100
50
0
0
100
200
Overall Survival (Months)

### Figure S6

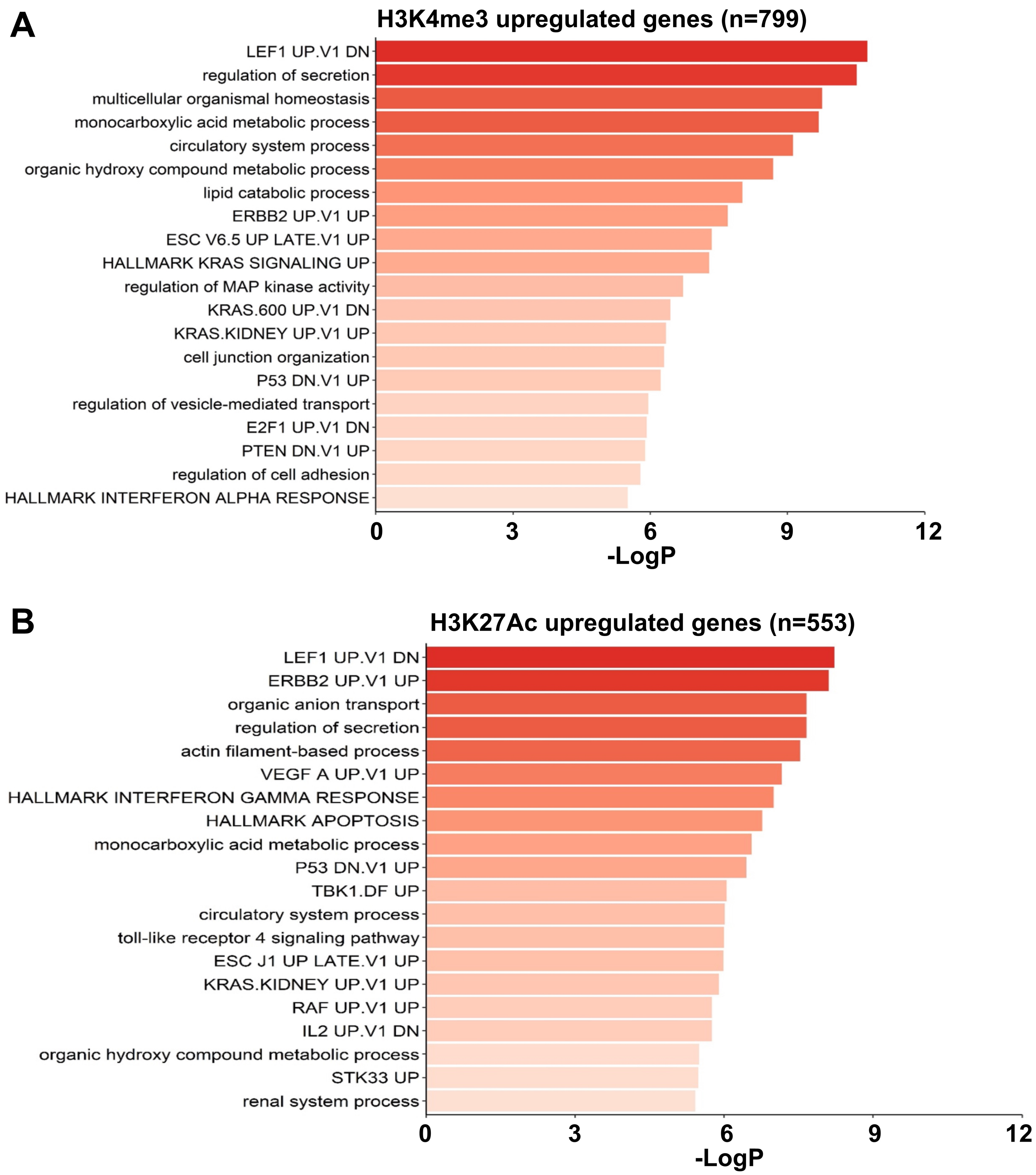
A
H3K4me3 upregulated genes (n=799)
LEF1 UP.V1 DN
regulation of secretion
multicellular organismal homeostasis
monocarboxylic acid metabolic process
circulatory system process
organic hydroxy compound metabolic process
lipid catabolic process
ERBB2 UP.V1 UP
ESC V6.5 UP LATE.V1 UP
HALLMARK KRAS SIGNALING UP
regulation of MAP kinase activity
KRAS.600 UP.V1 DN
KRAS.KIDNEY UP.V1 UP
cell junction organization
P53 DN.V1 UP
regulation of vesicle-mediated transport
E2F1 UP.V1 DN
PTEN DN.V1 UP
regulation of cell adhesion
HALLMARK INTERFERON ALPHA RESPONSE
0
3
6
9
12
-LogP
B
H3K27Ac upregulated genes (n=553)
LEF1 UP.V1 DN
ERBB2 UP.V1 UP
organic anion transport
regulation of secretion
actin filament-based process
VEGF A UP.V1 UP
HALLMARK INTERFERON GAMMA RESPONSE
HALLMARK APOPTOSIS
monocarboxylic acid metabolic process
P53 DN.V1 UP
TBK1.DF UP
circulatory system process
toll-like receptor 4 signaling pathway
ESC J1 UP LATE.V1 UP
KRAS.KIDNEY UP.V1 UP
RAF UP.V1 UP
IL2 UP.V1 DN
organic hydroxy compound metabolic process
STK33 UP
renal system process
0
3
6
9
12
-LogP

### Figure S7

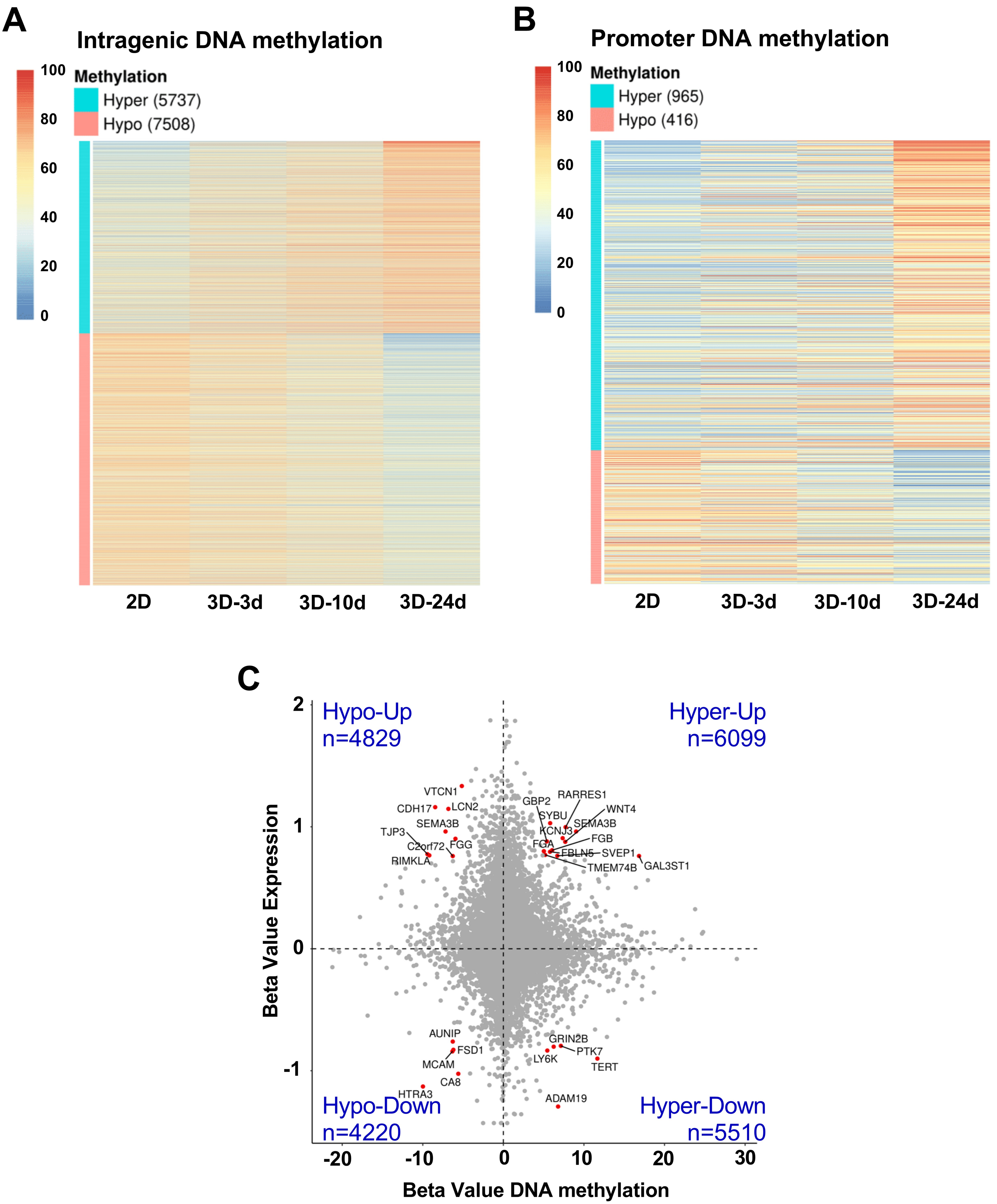
A
Intragenic DNA methylation
Methylation
Hyper (5737)
Hypo (7508)
100
80
60
40
20
0
2D
3D-3d
3D-10d
3D-24d
B
Promoter DNA methylation
Methylation
Hyper (965)
Hypo (416)
2D
3D-3d
3D-10d
3D-24d
C
Hypo-Up
n=4829
Hyper-Up
n=6099
Hypo-Down
n=4220
Hyper-Down
n=5510
Beta Value Expression
Beta Value DNA methylation
VTCN1
CDH17
LCN2
SEMA3B
TJP3
C2orf72
FGG
RIMKLA
GBP2
RARRES1
SYBU
WNT4
SEMA3B
KCNJ3
FGB
FGA
FBLN5
SVEP1
TMEM74B
GAL3ST1
AUNIP
FSD1
MCAM
CA8
HTRA3
GRIN2B
PTK7
LY6K
TERT
ADAM19

### Figure S8

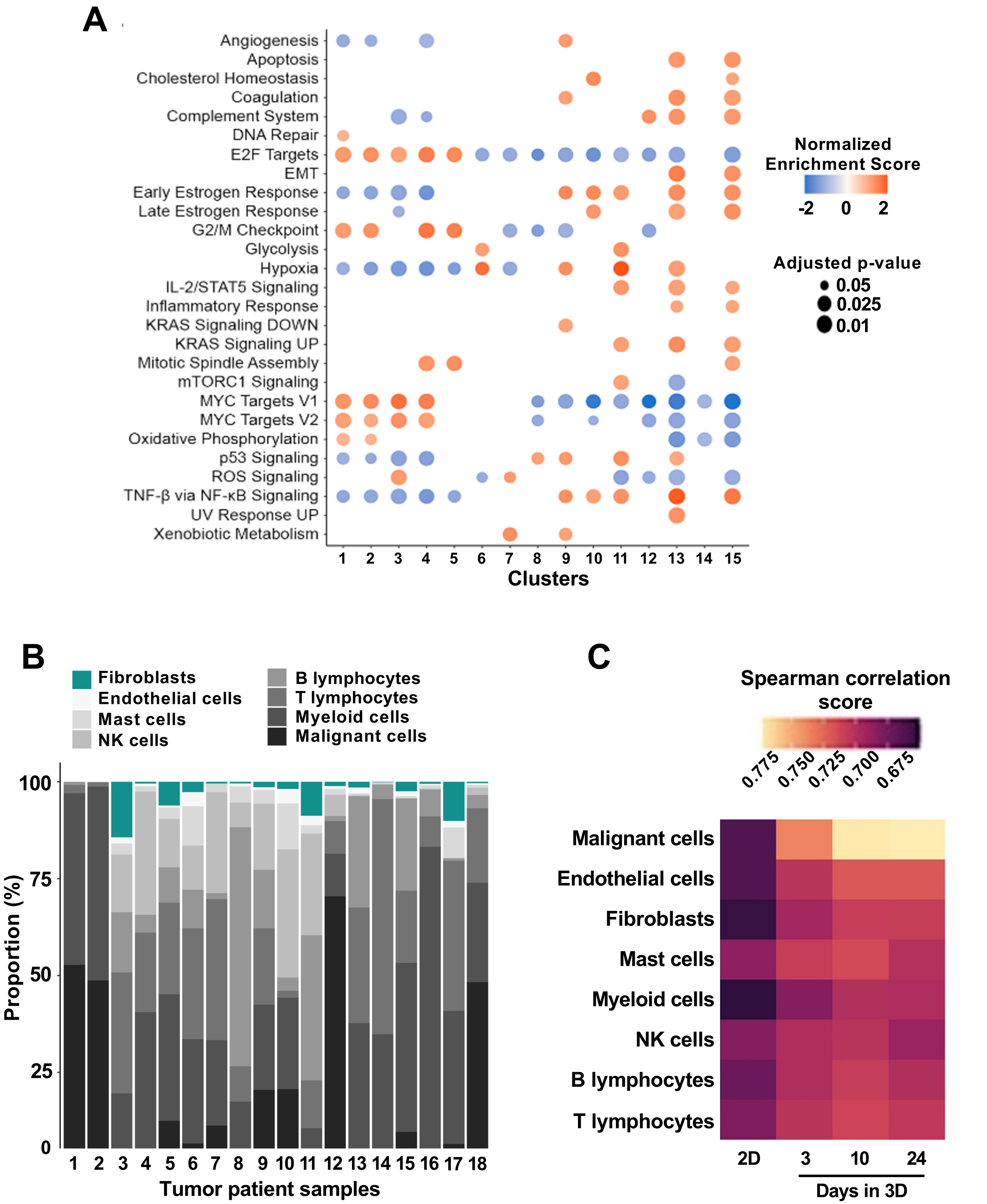
A
Angiogenesis
Apoptosis
Cholesterol Homeostasis
Coagulation
Complement System
DNA Repair
E2F Targets
EMT
Early Estrogen Response
Late Estrogen Response
G2/M Checkpoint
Glycolysis
Hypoxia
IL-2/STAT5 Signaling
Inflammatory Response
KRAS Signaling DOWN
KRAS Signaling UP
Mitotic Spindle Assembly
mTORC1 Signaling
MYC Targets V1
MYC Targets V2
Oxidative Phosphorylation
p53 Signaling
ROS Signaling
TNF-β via NF-κB Signaling
UV Response UP
Xenobiotic Metabolism
1 2 3 4 5 6 7 8 9 10 11 12 13 14 15
Clusters
Normalized Enrichment Score
-2 0 2
Adjusted p-value
0.05
0.025
0.01
B
Fibroblasts
Endothelial cells
Mast cells
NK cells
B lymphocytes
T lymphocytes
Myeloid cells
Malignant cells
Proportion (%)
0 25 50 75 100
1 2 3 4 5 6 7 8 9 10 11 12 13 14 15 16 17 18
Tumor patient samples
C
Spearman correlation score
0.775 0.750 0.725 0.700 0.675
Malignant cells
Endothelial cells
Fibroblasts
Mast cells
Myeloid cells
NK cells
B lymphocytes
T lymphocytes
2D 3 10 24
Days in 3D

### Figure S9

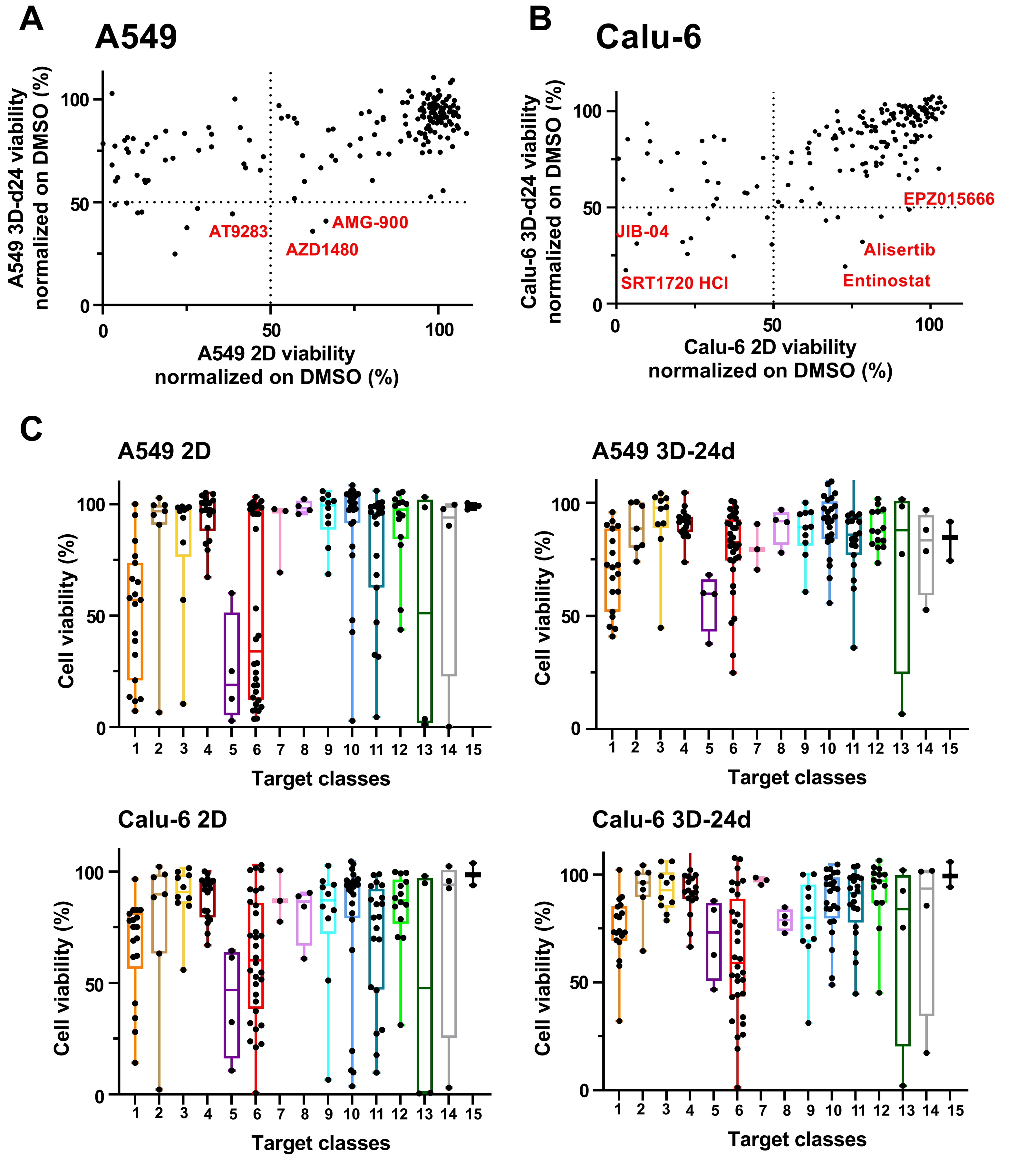


1. Aurora kinase
2. DNA methyltransferase
3. DNA/RNA synthesis
4. Epigenetic Reader Domain
5. FLT3
6. HDAC
7. HIF
8. Histone acetyltranferase
9. Histone demethylase
10. Histone methyltransferase
11. JAK
12. PARP
13. PIM
14. Sirtuin
15. Topoisomerase

### Figure S10

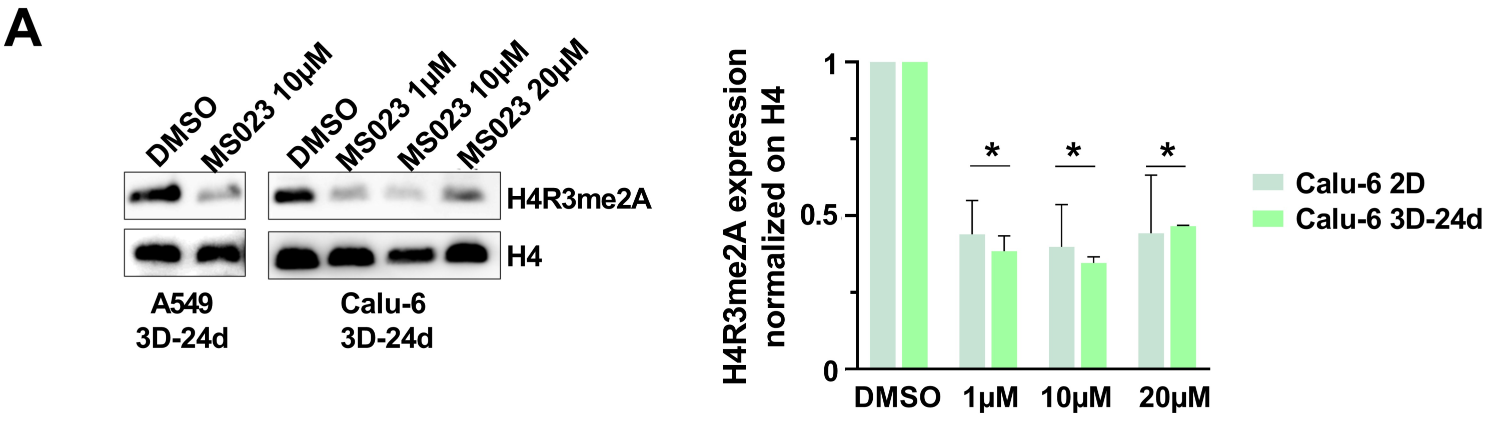
A
DMSO
MS023 10μM
DMSO
MS023 1μM
MS023 10μM
MS023 20μM
H4R3me2A
H4
A549
3D-24d
Calu-6
3D-24d
H4R3me2A expression normalized on H4
1
0.5
0
*
DMSO
1μM
10μM
20μM
Calu-6 2D
Calu-6 3D-24d

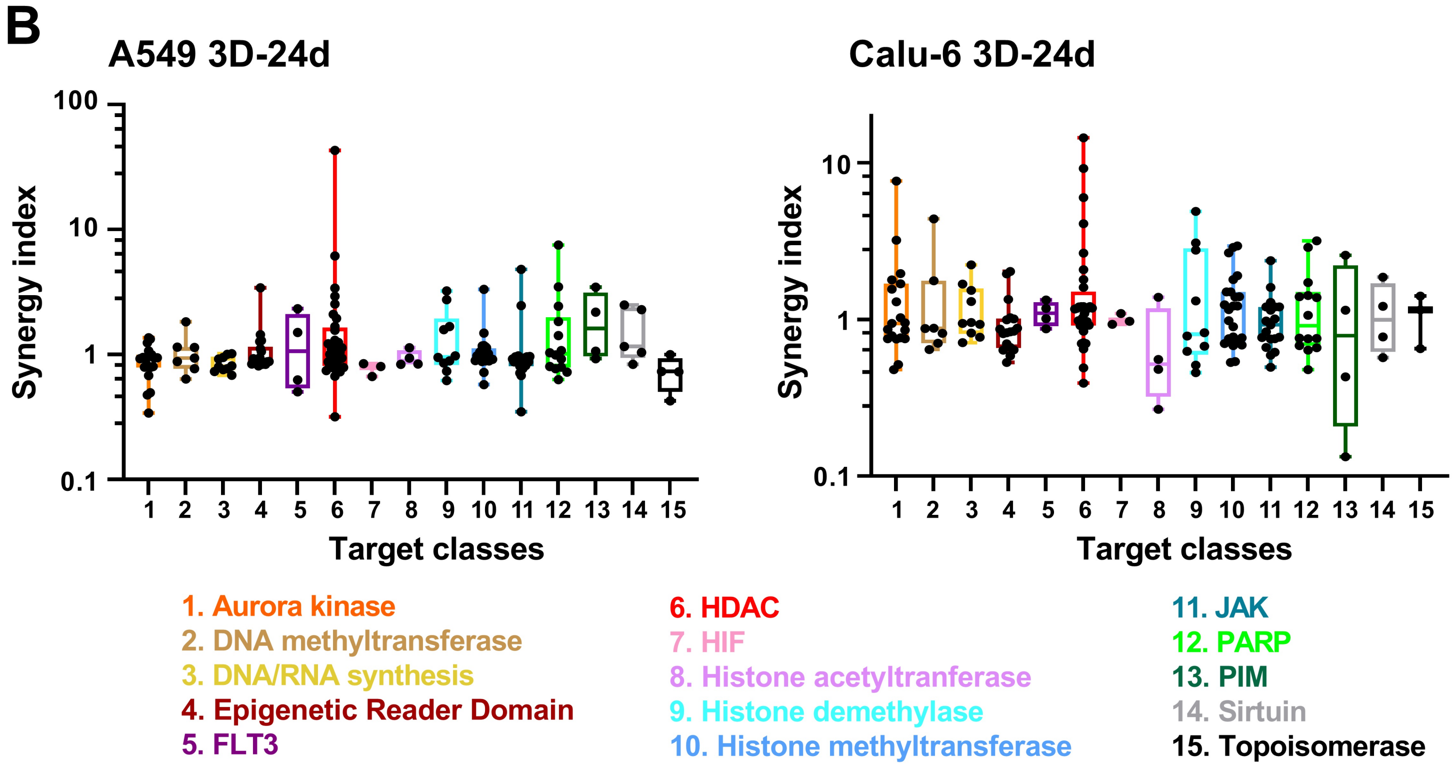
B
A549 3D-24d
Calu-6 3D-24d
Synergy index
100
10
1
0.1
1 2 3 4 5 6 7 8 9 10 11 12 13 14 15
Target classes
1. Aurora kinase
2. DNA methyltransferase
3. DNA/RNA synthesis
4. Epigenetic Reader Domain
5. FLT3
6. HDAC
7. HIF
8. Histone acetyltranferase
9. Histone demethylase
10. Histone methyltransferase
11. JAK
12. PARP
13. PIM
14. Sirtuin
15. Topoisomerase
